# Gene regulatory networks underlying human microglia maturation

**DOI:** 10.1101/2021.06.02.446636

**Authors:** Claudia Z. Han, Rick Z. Li, Emily Hansen, Hunter R. Bennett, Olivier Poirion, Justin Buchanan, Jean F. Challacombe, Bethany R. Fixsen, Samantha Trescott, Johannes C.M. Schlachetzki, Sebastian Preissl, Allen Wang, Carolyn O’Connor, Anna S. Warden, Shreya Shriram, Roy Kim, Celina T. Nguyen, Danielle M. Schafer, Gabriela Ramirez, Samuel A. Anavim, Avalon Johnson, Eniko Sajti, Mihir Gupta, Michael L. Levy, Sharona Ben-Haim, David D. Gonda, Louise Laurent, Christopher K. Glass, Nicole G. Coufal

## Abstract

The fetal period is a critical time for brain development, characterized by neurogenesis, neural migration, and synaptogenesis^1-3^. Microglia, the tissue resident macrophages of the brain, are observed as early as the fourth week of gestation^4^ and are thought to engage in a variety of processes essential for brain development and homeostasis^5-11^. Conversely, microglia phenotypes are highly regulated by the brain environment^12-14^. Mechanisms by which human brain development influences the maturation of microglia and microglia potential contribution to neurodevelopmental disorders remain poorly understood. Here, we performed transcriptomic analysis of human fetal and postnatal microglia and corresponding cortical tissue to define age-specific brain environmental factors that may drive microglia phenotypes. Comparative analysis of open chromatin profiles using bulk and single-cell methods in conjunction with a new computational approach that integrates epigenomic and single-cell RNA-seq data allowed decoding of cellular heterogeneity with inference of subtype- and development stage-specific transcriptional regulators. Interrogation of *in vivo* and *in vitro* iPSC-derived microglia models provides evidence for roles of putative instructive signals and downstream gene regulatory networks which establish human-specific fetal and postnatal microglia gene expression programs and potentially contribute to neurodevelopmental disorders.

## Main

To investigate mechanisms regulating human microglia maturation, we performed RNA-seq on FACS-sorted live microglia^14^ (Extended Data Fig.1a,b) isolated from early to mid-gestation fetuses (9W to 17W gestational age), which we refer to as fetal microglia, and from cortical tissue derived from epileptic resections of pediatric and adult patients (Supplementary Table 1), hereafter referred to as postnatal microglia. Additionally, we performed RNA-seq on a portion of fetal cortical tissue (i.e. whole fetal cortex) (Extended Data Fig.1c) to discern potential environmental cues that may drive fetal microglia phenotypes. Weighted gene correlation network analysis (WGCNA)^15^ identified 18 clusters of highly co-expressed gene modules with respect to both development and tissue type (Fig. 1a). Some modules were significantly enriched for genes essential to microglia and tissue macrophage function, such as leukocyte differentiation, migration, and phagocytosis (Supplementary Table 2). As expected, modules enriched for genes involved in axonogenesis and synapse organization were associated with fetal and postnatal cortical tissue. Modules associated with fetal, but not postnatal, microglia were enriched for genes with functional annotations for cell cycle and phagocytosis, suggesting that fetal microglia were more proliferative and phagocytic than postnatal microglia.

**Figure 1.**
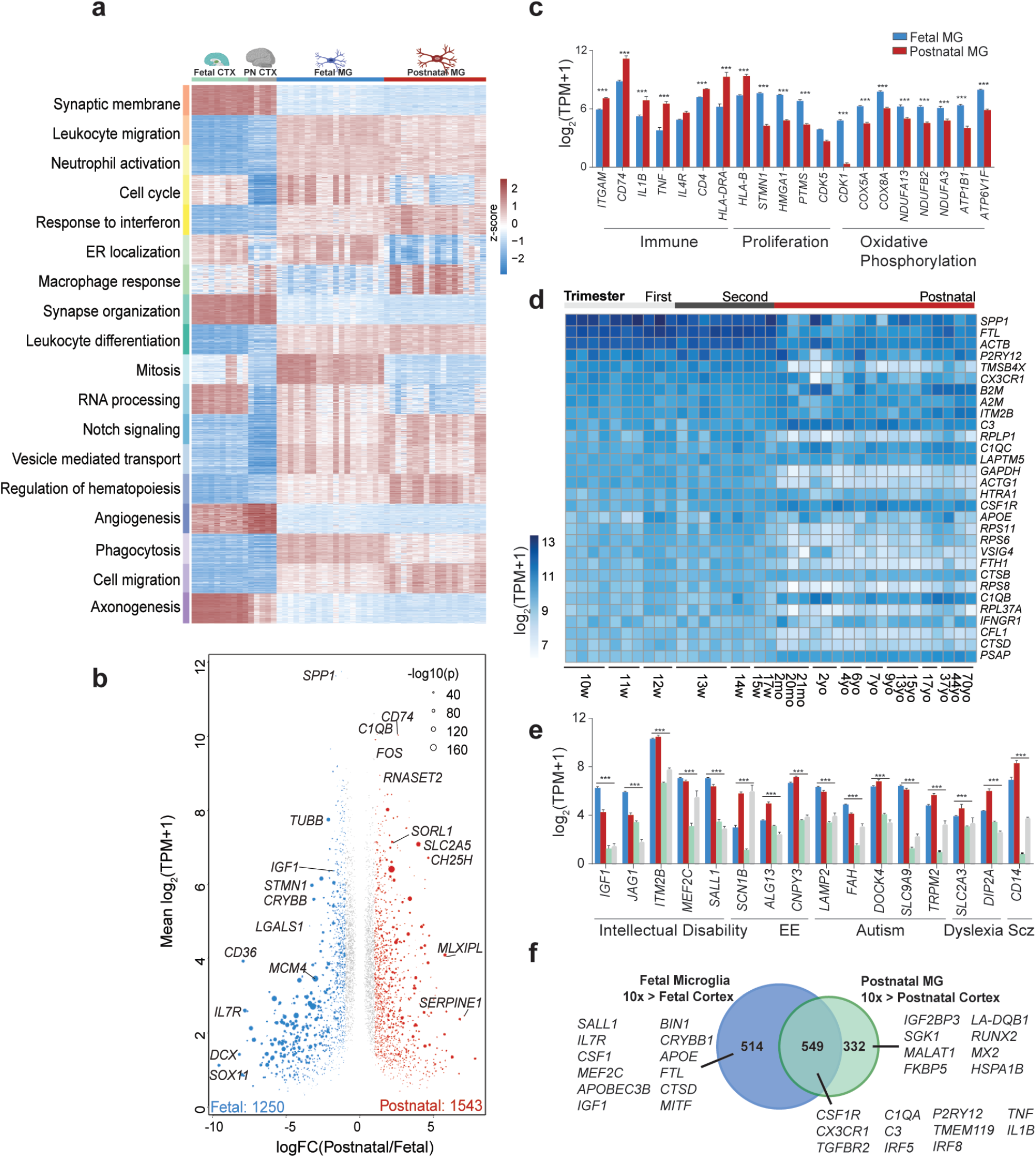
Human fetal microglia transcriptome. a. Heatmap expression z-scores of the top 100 genes from each significantly correlated module identified by WGCNA, ranked by Kleinberg’s hub centrality scores. b. MA plot of gene expression between human fetal (DEG, blue) and postnatal microglia (DEG, red). c. Bar charts of the expression levels of genes represented in GO term analysis of DEGs between fetal and postnatal microglia. d. Heatmap of gene expression representing the top 30 most variably expressed genes in fetal microglia. e. Bar charts of expression levels of monogenic NDD genes in microglia and brain cortex. f. Venn diagram illustrating overlap between genes enriched in fetal microglia as compared to fetal cortex (>10-fold, p-adj < 0.05 over bulk fetal cortex) and genes enriched in postnatal microglia compared to postnatal cortex12. Important genes related to microglia function are listed. *** indicates significant differential expression (> 2-fold, p-adj < 0.001).

Direct comparison of fetal and postnatal microglia transcripts identified approximately 3000 differentially expressed genes (DEGs, >2FC, p-adj <0.05, Fig. 1b) that clearly distinguished the two developmental stages. Gene ontology (GO) analyses of genes with higher expression in fetal microglia relative to postnatal microglia yielded strong enrichment for terms related to cell cycle, in agreement with the WGCNA analysis (Extended Data Fig. 1d). Fetal microglia expressed higher levels of genes related to oxidative phosphorylation (Fig. 1c), while postnatal microglia were enriched in genes associated with macrophage responses, including cytokine signaling and MHC protein complex (Fig. 1a, c). We also found significant individual variation in fetal and postnatal microglia, exemplified by the relative expression of the 30 most highly expressed genes in fetal microglia, including canonical microglia genes such as *CX3CR1* and *P2RY12* (Fig. 1d).

During early to mid-gestation, microglia play an essential role in modulating brain development. Maternal immune activation (MIA)^16^ has been linked to an increased risk of neuropathology, including autism spectrum disorder, with microglia posited to be one of the main mediators of MIA. We interrogated the differential expression of monogenic neurodevelopmental disorder (NDD) genes in microglia and whole cortex and across microglia development (Extended Data Fig.2a, b). We also included genes involved in monogenic mitochondrial disease and lysosomal storage disorders, as implicated by WGCNA and GO analyses. Numerous genes associated with intellectual disability, autism and schizophrenia are preferentially expressed in microglia compared to whole cortex (Fig. 1e, Extended Data Fig. 2a, b), suggesting a potential influential role for microglia in a variety of NDDs.

The microglia transcriptome is heavily influenced by local tissue environmental cues^12,17,18^. We previously identified 881 transcripts^12^ that were expressed >10-fold higher in human postnatal microglia than whole cortex at a false discovery rate of <0.05. Corresponding analysis of fetal microglia resulted in identification of 1063 genes that were expressed >10-fold higher in fetal microglia than fetal cortex, 549 of which were shared with the postnatal gene signature (Fig. 1f). As expected, shared genes include genes that specify microglia development and function, including *CSR1R, P2RY12* and cytokines such as *TNF* and *IL1β*. Notably, genes unique to the fetal microglial gene signature include those involved in maintaining microglia survival and proliferation, such as *CSF1* and *GAS6;* and adhesion and motility, including *LGASL1, ADGRE5, CAPG*, and *RAC2*. Alternatively, genes unique to the postnatal microglia gene signature are associated with immune response and myeloid activation, including *CD58, IRAK2*, and *MX2 (*Fig. 1f, Extended Data Fig. 3a)

### Environmental cues shaping microglia phenotypes

To identify potential environmental ligands and downstream signaling networks involved in brain development and/or fetal microglia function, we utilized NicheNet, which computationally predicts ligand-receptor interactions by combining gene expression with existing knowledge of signaling pathways and gene regulatory networks^19^. Top predicted ligand-receptor pairs and their target genes are shown in Fig. 2a. *IGF1*, a well-characterized growth factor important for neurogenesis, and Notch-ligand *DLL1* were two of the ligands predicted to preferentially influence the fetal microglia transcriptome, while *TGFβ1* and -3, *BMP7* were among the top predicted ligands for the postnatal period (Fig. 2b). Cytokines, including *IL6, TNF*, and *IL1β* were also predicted to affect postnatal microglia, paralleling the immune-related pathways identified by GO analysis.

**Figure 2.**
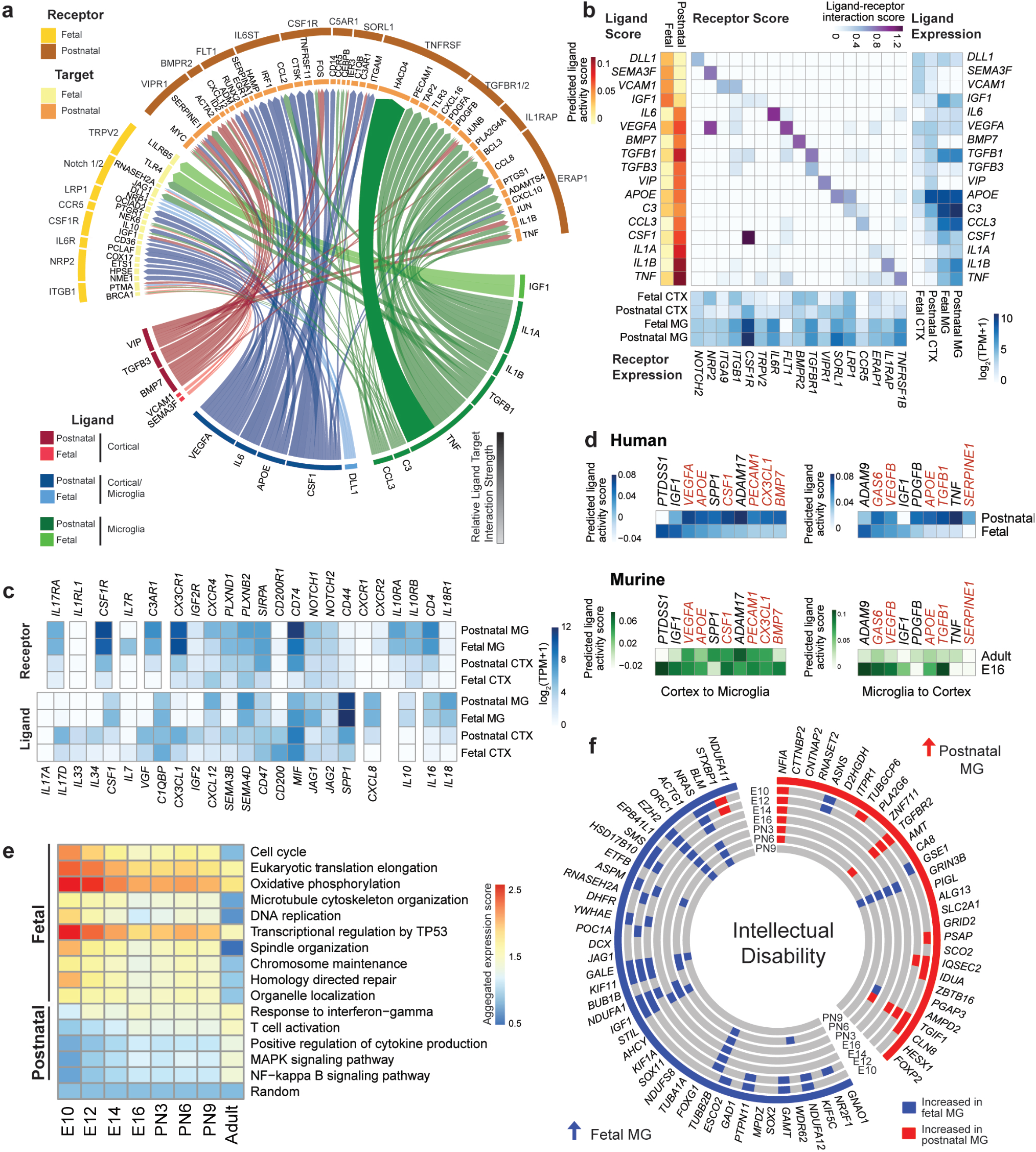
Gene expression in human and fetal microglia across development and disease. a. Circos plot indicating NicheNet prediction of ligand to target genes and predicted receptors in fetal and postnatal microglia. b. Heatmaps depicting ligand activity score (left) and ligand-receptor interaction score (middle) represented in 2a and of RNA expression of ligands (right) and receptors (bottom). c. Heatmap of gene expression of ligand (left)-receptor (right) pairs between human fetal bulk cortex, postnatal bulk cortex, fetal microglia and postnatal microglia. d. Heatmap showing NicheNet predicted ligands between human (blue, top) and mouse (green, bottom). Ligands are predicted for cortex signaling to microglia (left) and vice versa (right). Red labels ligands differentially predicted between murine and human. e. Heatmap showing aggregated expression scores (AES) of GO terms enriched in human microglia DEGs in murine microglia throughout development. f. Circos plot of monogenic NDD genes differentially expressed in human fetal (blue) compared to postnatal microglia (red) (outer most track). Each inner track shows whether the mouse ortholog is higher expressed in the indicated age compared to adult mouse microglia (blue) or vice versa (red). Grey denotes gene expression is nonsignificant between the indicated mouse microglia age and adult mouse microglia.

As the results from NicheNet are limited by the available content in its curated databases, we also manually evaluated the expression of additional ligand-receptor pairs. Interestingly, *CSF1* is preferentially expressed in fetal microglia, suggesting a critical autocrine role during the fetal period (Fig. 2c, Extended Data Fig. 3b). Additional ligands that may influence fetal microglia function include *JAG1*, a Notch ligand and *IL17D* (Fig.2c, Extended Data Fig.3c). Alternatively, postnatal microglia express higher levels of immune and migration-related ligands, including *SEMA4D, IL1β*, and *CXCL12* and higher levels of the signaling receptors *C3AR1, CXCR4, PLXNB2*, and *CD74 (*Fig.2c).

### Microglia developmental comparisons between mouse and human

While transcriptional networks driving human and mouse microglia identity are highly conserved, postnatal human and mouse microglia also exhibit significant differences in gene expression^12^. To investigate the evolution of these relationships from fetal to postnatal development, we utilized NicheNet to predict murine ligand-receptor pairs using previously published mouse microglia^20^ and embryonic whole cortex^21^ datasets. In examining paracrine signaling to microglia, *IGF1* and *PTDSS1* were predicted to influence human fetal microglia gene expression. In mice, *Apoe, Csf1*, and *Bmp7* were predicted to be mouse embryonic ligands; however, these ligands have higher predicted ligand activity in the human postnatal period as compared to the fetal period (Fig. 2d). Conversely, interrogation of potential ligand-receptor pairs involved in human and mouse microglia signaling to the brain parenchyma revealed *IGF1* as a top predicted ligand for embryonic/fetal microglia development and *TNF* and *PDGFB* as postnatal ligands (Fig. 2d).

We next calculated aggregated expression scores (AES) for the genes associated with GO terms enriched in human fetal and postnatal microglia (Extended Data Fig. 1g) and plotted these scores for mouse microglia throughout development. Embryonic mouse microglia at E10.5 were most similar to human fetal microglia, with higher AES associated with cell cycle, oxidative phosphorylation, and transcriptional regulation by TP53 (Fig. 2e). Conversely, there was a gradual increase in AES of human postnatal GO terms as mouse microglia matured (Fig. 2e).

Lastly, we examined the overlap in the expression of NDD monogenic genes between human fetal microglia and postnatal microglia and the orthologous comparison during mouse development. Lysosomal storage disorder genes are predominately expressed more highly in human postnatal microglia compared to fetal microglia. Conversely, monogenic mitochondrial disorder (Extended Data Fig. 3d) and intellectual disability genes (Fig. 2f) showed the opposite trend, with more genes being significantly higher expressed in fetal microglia versus postnatal microglia. Of the monogenic genes with orthologs in the mouse, there was little overlap in genes preferentially expressed during the corresponding development age between humans and mice (Extended Data Fig. 3d). For example, in intellectual disability, only a few genes, such as *NRAS, EZH2*, and *NFIA* show a similar pattern in preferential expression between fetal/mouse E10.5 and postnatal/adult mouse microglia (Fig. 2f). Collectively, these datasets suggest that the majority of microglia development between human and mice are conserved with some differences relating to age-specific expression of NDD genes.

### Differential chromatin accessibility between fetal and postnatal microglia

To infer roles of transcription factors (TFs) underlying microglia maturation, we defined regions of open chromatin in human fetal and postnatal microglia using bulk ATAC-seq. Comparison of distal and promoter elements between fetal and postnatal microglia delineated greater peak variability in distal elements (Fig. 3a) than in promoter proximal elements (Fig. 3b), as expected. *De novo* motif analysis of differential distal regions between human fetal and postnatal microglia revealed preferential enrichment of motifs assigned to MAF, MEF, CEBP, MITF/TFEB and NFY in fetal microglia (Fig. 3c) and to IRF and AP-1 in postnatal microglia (Fig. 3c). We then performed single cell ATAC-seq (scATAC-seq) on fetal and postnatal microglia to elucidate microglia subpopulation diversity. Clustering on scATAC-seq profiles revealed six distinct clusters, with segregation based on sample age (Fig. 3d). Cluster 0 contains contribution from all samples, while cluster 1,2, and 3 are biased towards fetal samples and cluster 4 and 5 are heavily represented by postnatal samples (Fig. 3e). Correlation analysis of all peaks between scATAC-seq samples and bulk ATAC-seq samples show high degree of similarity within respective postnatal and fetal groups (Extended Data Fig. 4a). Motif analysis of scATAC-seq using ChromVar^22^ confirmed enrichment for MAF, MAFB, TFE3 and TFEC motifs skewing towards the fetal samples, while motifs for AP1 and IRF factors were preferentially enriched in the postnatal samples (Fig. 3f-g, Extended Data Fig. 4b). As expected, the motif for SPI1, a macrophage lineage-determining factor, is present in all samples (Fig. 3g).

**Figure 3.**
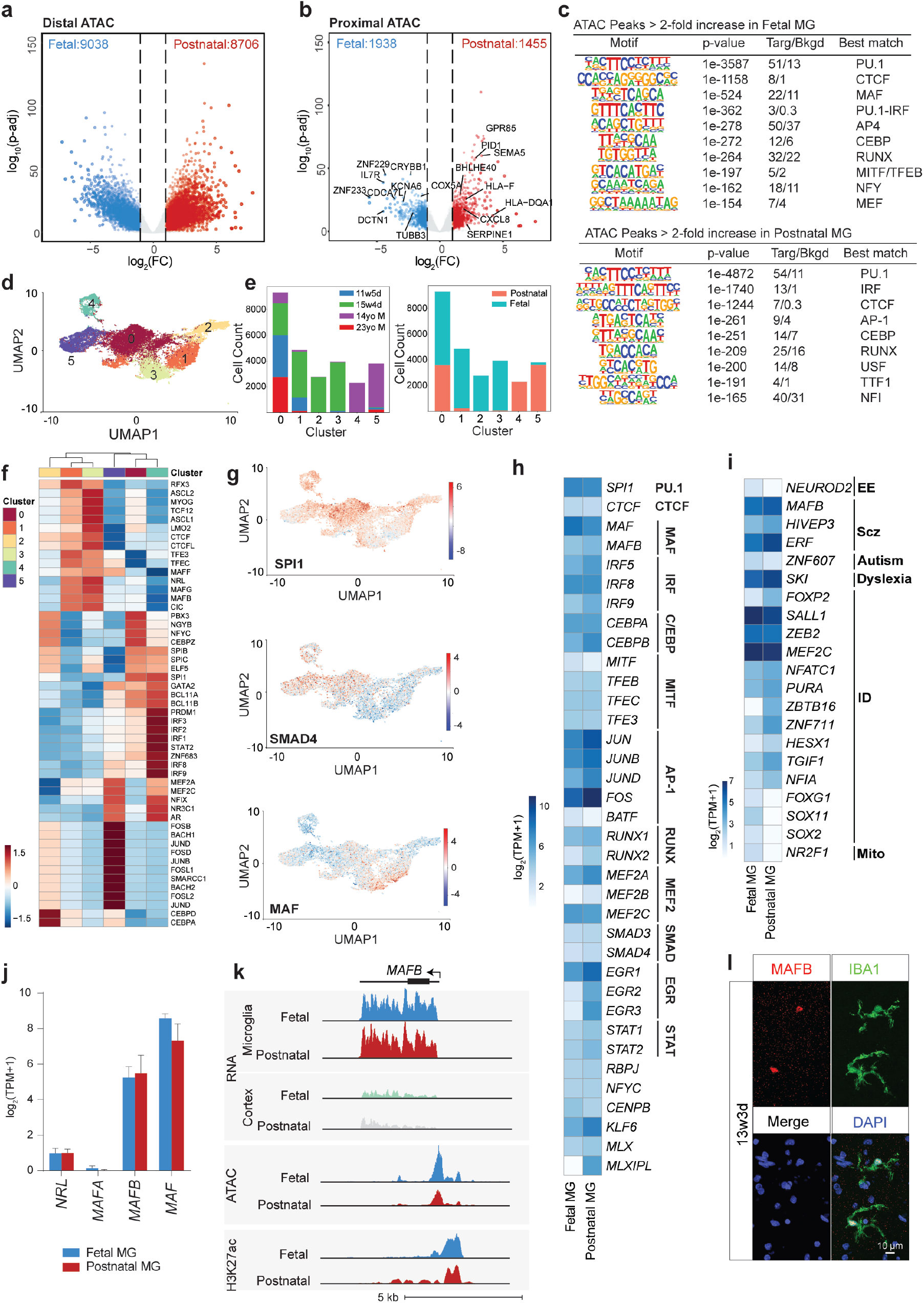
Fetal microglia have a distinct open chromatin profile and regulatory transcription factor network. a. Volcano plot of normalized bulk ATAC-seq tags at distal chromatin regions (>3kb from TSS) between human fetal and postnatal microglia. Differential regions enriched in fetal (blue) and postnatal (red) are colored. b. Volcano plot of normalized ATAC-seq tags at promoter proximal regions (<500bp from TSS) between human fetal and postnatal microglia. Differential regions enriched in fetal (blue) and postnatal (red) are colored. c. De novo motifs analysis of distal ATAC-seq peaks enriched in fetal (top) or postnatal (bottom) microglia. d. UMAP projection and color clustering of 27041 scATAC-seq profiles of fetal and postnatal microglia. Each dot represents one cell. e. Bar chart indicating sample contribution (left) and age contribution (right) to each cluster. f. Heatmap of average ChromVar score per motif and per cluster. Scores are averaged for all cells within each cluster and z-score normalized. g. UMAP visualization of enrichment for motifs associated with SPI1 (top), SMAD4 (middle) and MAF (bottom) using ChromVar. h. Heatmap of gene expression of transcription factors associated with motifs identified in bulk and scATAC-seq in human microglia. i. Heatmap of gene expression of monogenic NDD ranscription factors in human fetal and postnatal microglia. j. Bar chart of expression of selected MAF family members in human fetal and postnatal microglia. k. UCSC genome browser tracks of MAFB gene in human fetal and postnatal microglia and bulk cortex. l. Immunohistochemistry of fetal brain for MAFB and IBA1.

Expression levels of several TFs are correlated with the stage-specific motifs discovered in the bulk ATAC-seq and scATAC-seq analysis. For instance, selected AP-1 and IRF family members are more highly expressed in postnatal microglia (Fig. 3h), and as expected, MAF is preferentially expressed in fetal microglia. However, many of the differentially enriched motifs that are recognized by signal dependent TFs are primarily or strongly regulated at a post transcriptional level, including members of the AP-1, IRF, MITF/TFE, SMAD and MEF2 families. Altogether, our data suggests that differential enrichment for these motifs results from alterations in the brain environment between fetal and postnatal states. Additionally, a subset of TFs that are risk genes for NDDs are highly expressed in microglia and recognize motifs associated with open chromatin, including *MAFB* and *MEF2C* (Fig. 3f, i-k). Identification of high levels of *MAFB* expression in conjunction with a MAFB motif preferentially enriched in human fetal microglia was of interest because prior studies indicate that in the mouse, MAFB is exclusively expressed in postnatal microglia^20^. Immunostaining of MAFB in fetal cortical sections also show colocalization of nuclear MAFB with IBA1-positive microglia (Fig. 3l).

### Microglia Heterogeneity in development

To delineate microglial heterogeneity during human fetal development at the level of mRNA, we performed single-cell RNA-seq (scRNA-seq) on 5 fetal microglia and 3 postnatal microglia, yielding a total of 86,257 cells after quality control (Fig. 4a). Clustering with Harmony correction^23^ identified 10 clusters, with each cluster containing cellular contribution from every sample (Fig. 4b). RNA velocity^24^ analysis (Extended Data Fig.5a) identified strong intracluster movement in cluster 0 with a clear progression of fetal to postnatal microglia cells (Fig. 4c). This transition was associated with decreased expression of genes linked to endocytosis/phagocytosis and increased expression of genes involved in immune priming (Extended Data Fig. 5a, right). Expression of canonical microglia genes, such as *CX3CR1, CSF1R*, and complement proteins (e.g. *C1QA*) was relatively ubiquitous across cells (Fig. 4d, Extended Data Fig. 5b). However, some of these genes, such as *P2RY12* (Fig. 4d), and ligands such as *IGF1* (Extended Data Fig. 5c) show bias in expression towards fetal microglia.

**Figure 4.**
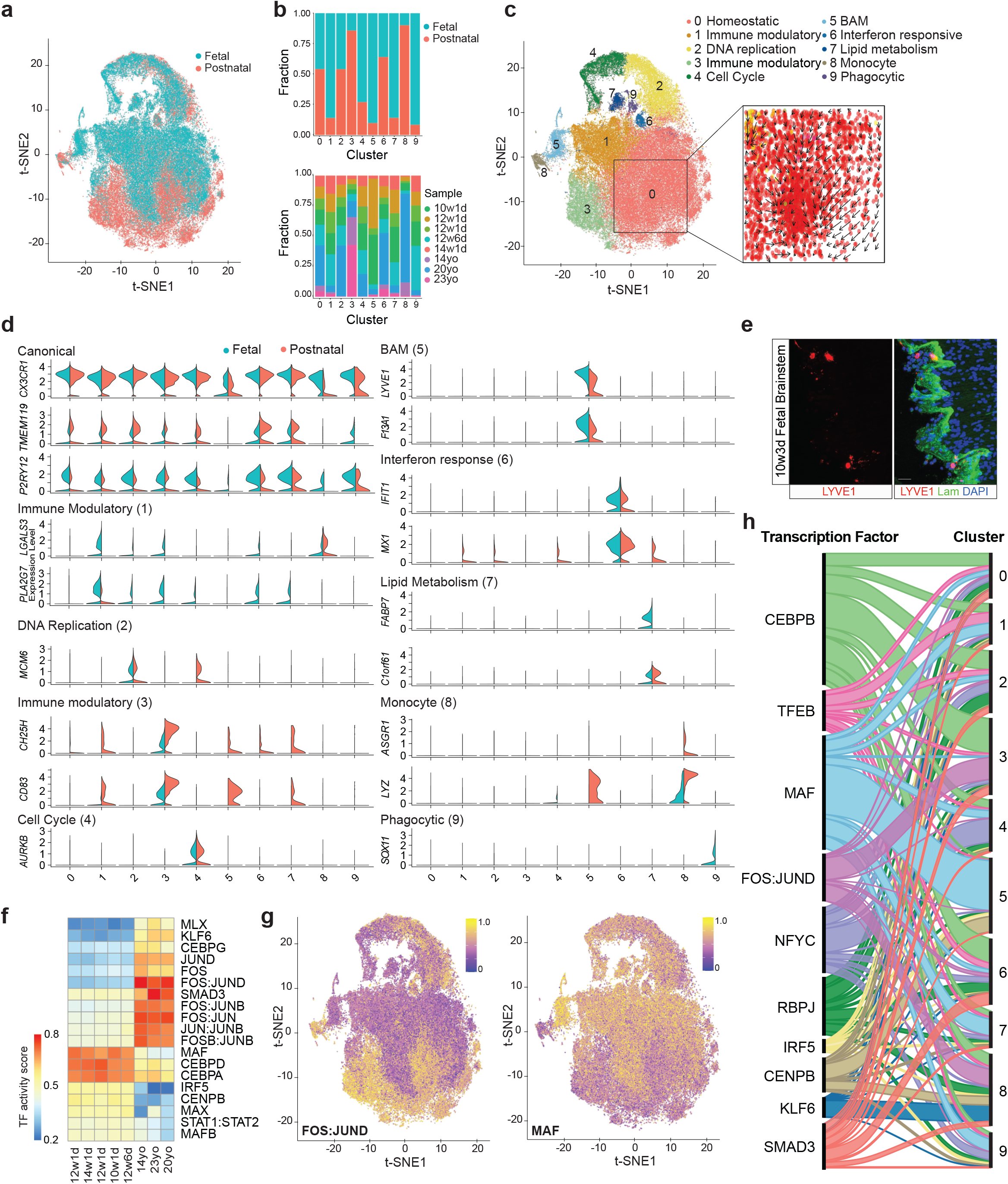
Single cell RNA-seq analysis reveals heterogeneity between fetal and postnatal microglial. a. tSNE projection of scRNA-seq analysis of fetal and postnatal microglia. Each dot represents one cell with coloring indicating age contribution. b. Bar graphs illustrating age contribution (top) and sample contribution (bottom) to each cluster. c. Annotation of scRNA-seq clusters with inset depicting RNA velocity analysis of cluster 0. d. Split violin plots showing the distribution of gene expression per cluster for fetal (salmon) and PN (teal). e. Immunohistochemistry of fetal brain for LYVE1 (BAM marker, red) and laminin (green). f. Heatmap depicting average TF activity scores for selected TFs, grouped by sample. g. tSNE projections of TF activity scores for FOS::JUND (left) and MAF (right). h. Alluvial plot representing TF activity scores per cluster with ribbon width proportional to TF activity score.

We detected two immune modulatory subgroups, cluster 1, expressing *LGALS3* and *UCP2*, and cluster 3, expressing *CD83, CH25H*, and *IL1B* (Fig. 4d, Extended Data Fig. 5d), with a fetal and a postnatal cell contribution bias, respectively. The postnatal immune modulatory microglia group has also been observed in other single cell analysis of human postnatal microglia^25^. Both fetal and postnatal microglia have subpopulations undergoing DNA replication (cluster 2) and cell division (cluster 4), characterized by expression of *MCM* genes (Fig. 4d), *E2F1*, and genes involved in microtubule dynamics, such as *STMN1, TUBB4B*, and *ASPM* (Fig. 4d, Extended Data Fig. 6a) and confirmed by immunofluorescence for division marker KI-67 (Extended Data Fig. 6b). In line with our bulk RNA-seq results, these two clusters had a statistically significantly higher contribution of fetal cells (57% cluster 2, 81% cluster 4, both p< 0.001 Chi square with Yates correction) (Extended Data Fig. 6c). We also detected a small interferon responsive cluster, cluster 6, expressing *IFIT1, IFIT3, MX1* (Fig. 4d, Extended Data Fig. 6d). There were several non-macrophage genes, such as *DCX* and *SOX11*, which mapped to a small cluster, cluster 9, that also co-expressed *CSF1R* (Extended Data Fig. 6e). Since these genes are strictly expressed in neural progenitor cells (NPCs), cluster 9 most likely captures microglia that have actively phagocytosed NPCs. Immunohistochemistry for the nuclear NPC marker SOX2 and cytoplasmic microglial marker IBA-1 identified rare events of likely phagocytosis (Extended Data Fig. 6f).

Lastly, we also detected a monocytic population (cluster 8), derived mostly from postnatal samples, and a border-associated macrophage (BAM) cluster (cluster 5), predominantly derived from fetal samples (Fig. 4d, Extended Data Fig. 7a). BAMs^26^ are composed of perivascular, meningeal, and choroid plexus macrophages and in mice, are phenotypically and transcriptionally distinct from parenchymal microglia^27,28^. Human BAMs have so far been uncharacterized in depth, with only perivascular macrophages described as CD163+. We used mouse BAM markers, such as *CD206*^*27-29*^ and *Lyve1*^*30*^, to qualify our human BAM clusters (Fig. 4d, Extended Data Fig. 7b). Immunostaining of fetal brain reveal LYVE1+ positive cells only near tissue borders, marked by laminin staining (Fig. 4e, Extended Data Fig. 7b,c).

### Enhancer-based inference of transcription factor activity from scRNA-seq

Although there has been extensive effort to utilize promoter sequences to assist the discovery of TFs driving specific cell identities^31,32^, enhancers have emerged as the dominant determinants of cell-type specific gene expression^33^. Thus, we integrated epigenomic and scRNA-seq data to infer the activities of enhancer-associated-TFs in microglia. Briefly, we identified enhancer sequences, using bulk ATAC-seq and H3K27acetylation ChIP-seq data, in fetal and postnatal samples and linked enhancers to target genes. Then we systematically identified TF motifs in the enhancer regions and computed a TF activity score for each enhancer-associated TF using scRNA-seq data, building a TF-gene network for each development age (Extended Data Fig. 8a). By this method, in general, TFs associated with highly expressed target genes are predicted to be more active than TFs associated with lowly expressed target genes.

TFs with higher activity scores in postnatal samples include KLF6, SMAD3, and AP-1 family members, while TFs with greater activity scores in fetal samples included STAT1::STAT2 dimer, IRF5 and developmental TFs such as MAF and MAFB (Fig. 4f,g). On a cluster-based analysis, cluster 3 is driven by CEBPB and FOS::JUND (Fig. 4h), consistent with this cluster being enriched for immune and primarily represented by postnatal microglia. The MiT/TFE family regulates transcriptional programming of autophagy and lysosome biogenesis^34^. Interestingly, TFEB, a MiT/TFE a family member, shows strongest activity for cluster 1 (Fig. 4h), the fetal-dominated immune modulatory cluster, correlating with the enrichment of lysosome-related genes including *CTSD, CD68*, and *LIPA* (Extended Data Fig. 5d). NFYC is most active in the cell cycle cluster, paralleling the function of NFY family members in regulating cell proliferation^35-37^. Clusters 1, 4, 5, 7 and 9 have greater MAF activity scores as compared to remaining clusters, most likely due to their higher composition of fetal cells compared to postnatal cells (Fig. 4h, Extended Data Fig.8b,c).

### iPSC-microglia in organoids and mouse brain capture distinct in vivo phenotypes

The recent ability to differentiate induced pluripotent stem cells (iPSCs) to microglia-like cells (iMGs), allows for functional studies *in vitro* and/or integration into cerebral organoids (oMGs) (Fig. 5a). Additionally, primitive hematopoietic progenitor cells (HPCs), also derived from iPSCs, can be transplanted into the brains of humanized immunodeficient mice, allowing for the development of human microglia that more closely resemble postnatal human microglia *in vivo*^38,39^ (Fig. 5a). Using Transcriptome Overlap Measure (TROM), a testing based method for identifying transcriptomic similarities^40^, we interrogated the similarity of human microglia, HPCs, iMGs, oMGs, and engrafted microglia (xMG)^38,39^. This analysis suggested that differentiation of human microglia in the mouse brain and introduction of iMGs into organoids capture distinct features of *ex vivo* fetal and postnatal microglia (Fig. 5c). Overall, comparison of the transcriptomes of oMGs to iMGs resulted in 528 DEGs (Extended Data Fig. 9a). GO analysis of DEGs expressed higher in oMGs compared to iMGs identified pathway enrichment in oMGs for cell cycle genes (Extended Data Fig. 9a,b). Interestingly, motif analysis of the distal accessible peaks differential in oMGs compared to iMGs (Extended Data Fig. 9c) show enrichment for motifs associated with AP-1, NRL, and TFE3 (Fig. 5d).

**Figure 5:**
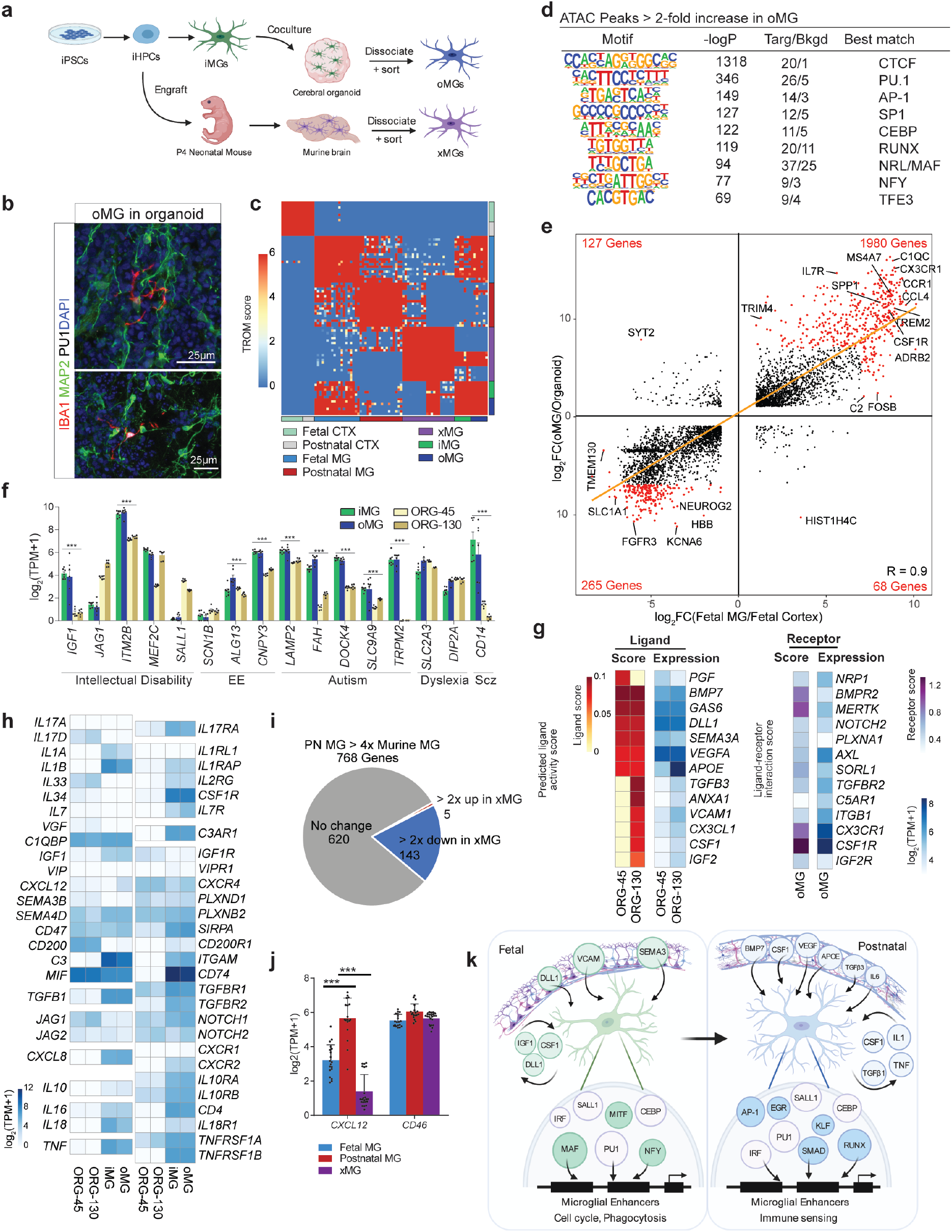
iPSC-derived microglia co-cultured with organoids is a suitable model for human fetal microglia. a. Schematic demonstrating derivations of HPCs, iMGs, oMGs, and xMGs from iPSCs. b. Immunohistochemistry depicting oMGs (IBA1, red; PU1, white) in proximity to neurons (MAP2, green) in organoids. c. TROM correspondence map of the transcriptomes of fetal and postnatal microglia and cortex, iMG, oMGs, and xMGs. Values are TROM scores with 0 being worst and 6 being best match. d. Motif enrichment analysis of distal differential accessible chromatin regions enriched in oMGs compared to iMGs. e. Ratio-ratio plot of genes comparing FC in gene expression between oMGs/cerebral organoids versus FC in gene expression between fetal microglia/fetal cortex. Pearson’s correlation coefficient is indicated in bottom right. f. Bar chart of expression of intellectual disability genes in iMGs, oMGs, and organoids aged day 45 and day 130. g. Heatmap depicting predicted ligand scores (left) in organoids aged day 45 and day 130, based on differentially expressed genes between fetal microglia and oMGs. RNA expression of ligands and receptors is shown on the right. h. Heatmap of gene expression of ligand (left)-receptor (right) pairs between iMGs, oMGs and organoids aged day 45 and day 130. i. Pie chart depicting the number of DEGs between adult mouse and human postnatal microglia that are expressed similarly between human postnatal microglia and transplanted microglia (xMGs) (grey), >2 FC in human postnatal microglia compared to xMGs (red), and > 2FC in xMGs compared to human postnatal microglia (blue). j. Bar chart of expression of selected genes in human fetal and postnatal microglia and xMGs. k. Schematic describing potential environmental cues and transcription regulatory networks that shape human fetal and postnatal microglia phenotypes. *** indicates significant differential expression (> 2-fold, p-adj < 0.001)

Thus we examined environmental similarities and differences between oMGs and human microglia. Comparison of the ratio of gene expression levels between oMG and organoids revealed a high degree of correlation to that of fetal microglia to fetal cortex, with r = 0.9 (Fig. 5e), suggesting that the organoid environment produces signals similar to the fetal brain. Notably, *SALL1*, a transcriptional regulator of microglia homeostasis *in vivo*, is not induced in iMGs in the organoid environment, suggesting that brain derived signals necessary for its expression are not produced in this system (Fig. 5f). Expression levels of NDD genes, including those associated with autism, intellectual disability (Fig. 5f) and lysosomal storage disorders in oMGs and organoids closely mirrored that of primary fetal brain derived cells (Extended Data Fig. 10a). Similar to fetal microglia, oMGs displayed transcriptomic evidence of NPC phagocytosis (Extended Data Fig. 10b).

NicheNet analysis predicted organoid ligands that may drive differentially higher expressed genes in oMGs compared to fetal microglia, including *BMP7, VEGFA*, and *TGFβ3* (Fig. 5g). Interestingly, these ligands were predicted to have greater ligand activity during the human postnatal period, suggesting that the organoids also capture components of postnatal environmental cues. Probing of other ligand-receptor pairs reveals that there is a high degree of concordance in expression of receptor-ligand pairs in oMGs/iMGs compared to human microglia, including cytokines, such as *IL10, IL1A/B*, growth factors like *IGF1*, and essential factors for microglia survival and identity such as *TGFB1, CSF1, and IL34* (Fig. 5h, Extended Data Fig. 10c).

In contrast to organoids, engraftment of microglia into the mouse brain results in upregulation of *SALL1* and a gene expression program more resembling *ex vivo* microglia than *in vitro* microglia^38^. However, significant differences remain, which could be due to differences between the mouse and human brain environment. To address this possibility, we investigated the expression levels of 768 genes that are expressed >4-fold more highly in human microglia compared to mouse microglia^12^ in the xMG transplant model (Fig. 5i). We found that 620 of these human-specific genes retained a human-specific pattern of expression in the context of the mouse brain, being approximately equal to levels observed in postnatal microglia, exemplified by *CD46* (Fig. 5j). These results are consistent with human-specific expression being due to divergence in *cis*-active regulatory elements rather than differences in brain environment and support the use of the explant model for studies in which these genes are relevant. Conversely, 143 of the human-specific genes exhibited a greater than 2-fold reduction in expression in xMGs, exemplified by *Cxcl12* (Fig. 5j). These results are consistent with either reduced levels or cross species incompatibilities of important signaling molecules. Further, they provide an indirect validation of the NicheNet analyses, which predicted *CXCL12* to be a downstream target gene of *BMP7* signaling in both the postnatal human cortex and organoid model, but not the mouse cortex.

## Discussion

Analyses of the transcriptomes of human fetal and postnatal microglia in relation to the surrounding cortex reveals substantial microglia maturation and indicates that a broad spectrum of genes associated with monogenic neurodevelopmental disorders are preferentially expressed in microglia. Many of these genes exhibit significantly different patterns of expression in mouse microglia. Integration of transcriptomic and epigenetic data from human samples and recently developed iPSC-dependent model systems confirm previously established environmental factors and enable inference of roles of numerous additional ligands, receptors and transcription factors in regulation of human microglia phenotypes. Enhanced proliferative and phagocytic phenotypes of fetal microglia are associated with cell autonomous expression of *CSF1, IGF1* and *DLL1* and evidence for preferential activities of MAF, NFY and MITF/TFE families of transcription factors, respectively (Fig. 5k). Microglia maturation during the transition to the postnatal state is characterized by acquisition of an immune competent phenotype driven by cortically derived *CSF1, VEGF, BMP7, APOE, TGFβ3* and cell autonomous production of *IL1β, TGFβ, TNF* and *CSF1*. These factors are proposed to preferentially regulate the activities of AP-1, SMAD, KLF and EGR family members (Fig. 5k). Organoid and *in vivo* engraftment systems capture distinct and complementary aspects of human microglia maturation and thus have the potential to provide powerful model systems for further understanding mechanisms underlying human-specific programs of gene expression.

## Methods

### Human tissue

Microglia were isolated from postnatal brain tissue (in excess of that needed for pathological diagnosis) as previously described^1^. All postnatal patients were diagnosed with refractory epilepsy and had epileptogenic focus resections at either Rady Children’s Hospital or through the UC San Diego Medical System (Jacobs Medical Center or UC San Diego Hillcrest Hospital). Brain tissue was obtained with informed consent from adult patients, or by informed parental consent and assent when applicable from pediatric patients under a protocol approved by the UC San Diego and Rady Children’s Hospital Institutional Review Board (IRB 160531, IRB 171361). Resected brain tissue was immediately placed on ice and transferred to the laboratory for microglia isolation or post fixation for histology within three hours after resection. Charts were reviewed for final pathological diagnosis, epilepsy medications, demographics, and timing of stereoelectroencephalography (SEEG) prior to surgery. Fetal brain samples were collected under a protocol approved by the UC San Diego Institutional Review Board (IRB 171379). Fetal brain samples were obtained within 1 hour of the pregnancy termination procedure after informed consent and transported in saline then were immediately either utilized for microglial isolation or were postfixed for histology. The reported data sets are from sequential samples for which cell viability and sequencing libraries met technical quality standards. No other criteria were used to include or exclude samples. All relevant ethical regulations were complied with.

### Human microglia isolation

Human brain tissues were manually dissected into small 2-3 mm^3^ pieces and immersed in homogenization buffer (HBSS (Life Technologies, 14175-095), 1% bovine serum albumin (Sigma-Aldrich, A3059), 1 mM EDTA)) for mechanical dissociation using a 2 ml polytetrafluoroethylene pestle (Wheaton, 358026)^1^. Postnatal human microglia were isolated using an approach that combined Percoll enrichment and flow cytometry purification. Brain homogenate was pelleted, filtered through 40μm filter, re-suspended in 37% isotonic Percoll (Sigma, P4937) and centrifuged at 600xg for 30 min at 16-18°C with minimal acceleration and no deceleration. Percoll gradients were utilized for all postnatal samples and only for fetal samples > 500mg. Following Percoll gradient centrifugation, pelleted cells were collected and washed twice with homogenization buffer, filtered with a 40 µm strainer (BD Falcon 352350) and incubated with Fc-receptor blocking antibody (Human TruStain FcX, BioLegend 422302) in homogenization buffer for 20 minutes on ice. Then cells were stained with the following cell surface marker antibodies for 30 min on ice (1:100 dilution, all from BioLegend): CD11b-PE (301306, clone ICRF44,), CD45-APC/Cy7 (304014, clone HI30), CD64-APC (305014, clone 10.1), CX3CR1-PerCP/Cy5.5 (341614, clone 2A9-1), CD14-AF 488 (301811, clone M5E2), HLA-DR-PE/Cy7 (307616, clone L243), and CD192-BV510 (357217, clone K036C2). CD14 and HLA-DR were included to further characterize immune cells but did not further discriminate subsets of microglia. Zombie Violet (Biolegend, 423113) or DAPI was added to the samples for viability discrimination immediately prior to sorting (1 µg/ml final concentration). Microglia were purified with either a BD Influx (100-µm nozzle, 22 PSI, 2-drop purity mode, sample chilling) or BD FACS AriaFusion (100-µm nozzle, 20 PSI, Purity mode (a 1-2 drop sort mode), sample chilling) and defined as live/DAPI^-^/Zombie violet^-^ CD11b^+^CD45^Low^CD64^+^CX3CR1^High^ single cells (Fig. S1A). Flow cytometry data were also analyzed using FlowJo software (Tree Star).

### Human pluripotent stem cell culture

All studies were conducted according to the human stem cell (hESCRO) protocol approved by the Embryonic Stem Cell Research Oversight (ESCRO) Committee at University of California, San Diego (IRB 171379). Human embryonic stem cell (ESC) line H1 (WiCell Research Institute, Madison, WI) ^2^ and induced pluripotent stem cell (iPSC) line EC11, derived from primary human umbilical vein endothelial cells (Lonza, Bioscience) ^3^, and an additional iPSC cell line derived from a control human aging cohort (UKERfG3G-X-001) ^4^, were cultured utilizing standard techniques. In brief, cells were cultured in StemMacs iPS-Brew media (Miltenyi Biotech, Auburn, CA) and routinely passaged utilizing Gentle Cell Dissociation Reagent (STEMCELL Technologies) onto Matrigel-coated (1 mg ml^-1^) plates. Karyotype was established by standard commercial karyotyping (WiCell Research Institute, Madison, WI).

### Microglial Differentiation

Microglia were generated as previously described with minor modification^5^. Briefly, ESC/iPSCs were plated in iPS-Brew with 10 μM ROCK inhibitor (Stem Cell Technologies) onto Matrigel-coated (1 mg ml^-1^) 6-well plates ^6^ using ReLeSR (STEMCELL Technologies). Cells were differentiated to CD43+ hematopoietic progenitors using the StemCell Technologies Hematopoietic Kit (Cat #05310). On day 1, cells were changed to basal media with supplement A (1:200), supplemented with an additional 1 ml/well on day 3, and changed to basal media with supplement B (1:200) on day 3-4 depending on cellular morphology. Cells received an additional 1 ml/well of medium B on days 5, 7, and 10. Nonadherent hematopoietic cells were collected between days 11-14 depending on the differentiation. Cells were then replated onto Matrigel-coated plates (1 mg ml^-1^) at a density of 300,000 cells/well in microglia media. Microglia media consisted of DMEM/F12 (Thermofisher), 2x insulin-transferrin-selenite (Gibco), 2x B27 (Lifetech), 0.5x N2 (Lifetech), 1x GlutaMAX (Gibco), 2x non-essential amino acids (Gibco), 400 μM monothioglycerol, and 5 μg ml^-1^ insulin (Sigma). Microglia media was supplemented with 100 ng ml^-1^ IL-34 (Proteintech), 50 ng ml^-1^ TGFβ1 (Proteintech) and 25 ng ml^-1^ M-CSF (Proteintech). Cells were supplemented with microglia media with IL-34, TGFβ1 and M-CSF every other day. 25 days after initiation with microglia media, cells were resuspended in microglia media with IL-34, MCSF and TGFβ1 with the addition of CD200 100 ng ml^-1^ (Novoprotein) and CX3CL1 100 ng ml^-1^ (Peprotech). Cells were collected on Day 28 for experiments.

### Organoid Differentiation

Cerebral organoids were generated as previously described ^7^ with modifications. Briefly, iPSCS were grown on Matrigel, then washed with DMEM/F12 (Sigma) and were dissociated with collagenase at a concentration of 1.5mg/mL for 1 hour until colonies floated from the plate. Colonies were washed with DMEM/F12 several times and allowed to settle by gravity then were plated in embryoid body formation media (DMEM/F12 supplemented with 2mM GlutaMAX, 1% non-essential amino acids (Thermofisher), 50nM 2-mercaptoethanol (Gibco), 20% knockout serum replacement, 2 µm dorsomorphin and 2 µm A83) with the addition of ROCK inhibitor Y27632 (40 µM) and 50ng mL^-1^ of bFGF-2 in an ultralow attachment plate (Day 1). On days 3 and 5 cells were fed with EB formation inhibitor without ROCK inhibitor or FGF. On day 5 organoids were transitioned to media for neural induction, consisting of DMEM/F12 with 2mM GlutaMAX, 1x N2 supplement (Gibco), 1% NEAA, 10ug mL^-1^ Heparin (Sigma), 1mM CHIR99201 and 1mM SB431542. On day 7 embryoid bodies were manually embedded in 18μL droplet of Matrigel for 30min at 37°C. Organoids were transitioned from neural induction to long term differentiation media on Day 12-14, consisting of DMEM/F12 with 2mM GlutaMAX, 1x N2, 1x B27 (Thermofisher), 2.5ug ml^-1^ insulin, 55 µM 2-mercaptoethanol, 1% NEAA, 1% pen/strep. At this time organoids were moved to an orbital shaker for the remainder of the culture time. Excess Matrigel was manually removed on day 20, and cerebral organoids were utilized at 8-12 weeks of age for delineated studies.

### Immunofluorescence staining and analysis of cerebral organoids

Organoids were fixed in 4% paraformaldehyde in 0.1 M Phosphate buffer saline for 45 - 60 minutes at 4°C and washed three times in PBS, then cryoprotected in 30% sucrose and embedded in tissue freezing medium (GeneralData) for cryo-sectioning. Twenty-μm sections were cut on a cryostat, mounted on Superfrost plus slides (Thermo Scientific, Menzel-Glaser), and stored at -80°C until staining. For immunofluorescence, sections were rehydrated, rinsed in 0.1 M TBS, then permeabilized and blocked for non-specific binding in blocking buffer containing 3% normal horse serum and 0.25% Triton X-100 (Sigma X100) in a humidified chamber for 1 hr at room temperature^8^. Slides were then incubated with the appropriate primary antibodies diluted in blocking buffer at 4°C overnight. The next day, sections were washed twice (fifteen minutes each) in 0.1M TBS, washed with blocking buffer (once for 30 minutes), and incubated with fluorophore-conjugated secondary antibodies diluted in blocking solution at RT for 2 hrs. After the two-hour incubation, sections were counter stained with DAPI for 10 minutes, rinsed three times in 0.1M TBS (15 minutes each), rinsed with 0.1 M PO_4_, and mounted with Shandon Immuno-Mount (Thermo Scientific, 9990412). Imaging was performed on a Leica TCS SPE confocal microscope or a Nikon Eclipse Ti2-E with laser scanning confocal A1R HD.

### Isolation of iMGs and oMGs

For iMGs, cells were carefully manually scraped from the Matrigel coated plate and were concentrated by a 300 rcf x 5 minutes. For oMGs, organoids were carefully collected, allowed to settle by gravity, and then dissociated mechanically in staining buffer (HBSS 1x with 1mM EDTA and 1% BSA) using a 2 ml polytetrafluoroethylene pestle (Wheaton, 358026) in a fashion identical to fetal brain tissue. No percoll gradient was utilized.

Both iMGs and oMGs were resuspended in staining buffer and were blocked with Fc receptor blocking antibody (Human TruStain FcX, BioLegend 422302) for 10 minutes. Both iMG and oMGs were stained with the following 6 antibodies, all at 1:30 dilution and all from Biolegend: CD64-APC (305014), CX3CR1 PCP-Cy5.5 (341614), CD14-488 (325610), CD11b-PE (301309), HLADR PE-Cy7 (307616), CD45-APC-CY7 (368516) for one hour. Cells were then washed and incubated in Zombie Violet (1:1000, Biolegend) for live/dead discrimination. Controls consisted of cells incubated with a combination of appropriate isotypes for each antibody (Biolegend). Microglia were purified on a BD InFlux Cytometer (Becton-Dickinson).

### Tissue processing for immunostaining

For fixation, fetal and postnatal tissue was fixed in 4% formaldehyde in phosphate-buffered saline (PBS) overnight at 4°C then transferred to 30% sucrose. The tissue was sectioned in 20-μm sections using a cryostat. Sections were stored at -80°C until staining. For IL17 receptor staining, heat-induced antigen retrieval was performed using (standard protocols) Target Retrieval Solution (Dako S1699). Target Retrieval Solution was diluted to 1x concentration and then heated to 90°C. Tissue was incubated in solution at 90°C for 30 minutes followed by three washes in 0.1M TBS (Trizma Base (Sigma T1503), NaCl (Sigma S9888), KCl (Sigma P9541), HCl (Sigma H1758), pH 7.5). After the washes staining was continued per the immunostaining protocol below.

### Immunostaining

Sections were washed in 0.1M TBS and then non-specific binding site blocking and cell permeabilization was performed with blocking buffer containing 3% normal horse serum and 0.25% Triton X-100 (Sigma X100). Sections were incubated with primary antibody (see list below) in blocking buffer at 4°C overnight. After washing in 0.1M TBS, sections were incubated with secondary antibodies (see list below) for two hours at room temperature (1:250, Jackson Laboratories). Sections were washed with TBS before nuclear counterstaining (DAPI 1:1000, Thermo Fisher) and then mounted with Shandon Immu-Mount (Thermo Scientific, 9990412). Imaging was performed on a Leica TCS SPE confocal microscope or a Nikon Eclipse Ti2-E with laser scanning confocal A1R HD.

Cell cultures were fixed with 4% paraformaldehyde solution for 30 minutes at room temperature. Antigen blocking and cell permeabilization were performed using blocking buffer consisting of 3% horse serum and 0.25% Triton X-100 (Sigma) in TBS for 1 hour at room temperature. Primary antibodies were incubated in 3% horse serum overnight at 4°C, and secondary antibodies (1:250, Jackson Laboratories) were incubated in the same solution for 1 hour at room temperature. The cells were counterstained with DAPI for nuclei detection and mounted with Shandon Immuno-Mount (Thermo Scientific, 9990412).

The following primary antibodies were used: goat Iba1 (1:200, Abcam, ab5076), rabbit Iba1 (1:500, Wako, 019-19741), mouse Nestin (1:500, EMD Millipore, ABD69), chicken MAP2 (1:500, Abcam, ab5392), rabbit PU1 (1:250, Cell Signaling, 2266S), rabbit P2RY12 (1:200, HPA014518), rat CX3CR1 (1:100, Biolegend, 341602), goat IL17 (1:100, R&D Systems, AF-317-NA), mouse IL17R (1:100, R&D Systems, MAB177), rabbit Sox2 (1:200, Cell Signaling, 2748S), rabbit MafB (1:200, Abcam, ab223744), mouse Ki67 (1:1000, Leica, ACK02), rabbit Lyve1 (1:200, Abcam, ab36993), mouse CD163 (1:100, BIORAD, MCA1853), rabbit Laminin (1:100, EMD Millipore, MABE622), mouse Laminin (1:100, Telios/Gibco BRL).

The following secondary antibodies were used (all at 1:250): Donkey Cy3 anti Goat (Jackson Laboratories, 705-165-147), Donkey Alexa Fluor 488 anti-Goat (Jackson Laboratories, 705-545-147), Donkey Alexa Fluor 647 anti-Goat (Jackson Laboratories, 705-175-147), Donkey Cy3 anti Rabbit (Jackson Laboratories, 711-165-152), Donkey Alexa Fluor 488 anti-Rabbit (Jackson Laboratories, 711-545-152), Donkey Alexa Fluor 647 anti-Rabbit (Jackson Laboratories, 711-175-152), Donkey Cy3 anti Mouse (Jackson Laboratories, 715-165-151), Goat Alexa Fluor 488 anti-Mouse (Jackson Laboratories,715-545-151), Donkey Alexa Fluor 647 anti-Mouse (Jackson Laboratories, 715-545-151), Donkey Alexa Fluor 488 anti-Chicken (Jackson Laboratories, 703-545-155), Donkey Alexa Fluor 647 anti-Rat (Jackson Laboratories, 712-605-153).

### Assay for Transposase-Accessible Chromatin-sequencing (ATAC-seq)

30,000-50,000 isolated human microglia, iMGs or oMGs were lysed in 50 µl lysis buffer (10 mM Tris-HCl pH 7.5, 10 mM NaCl, 3 mM MgCl_2_, 0.1% IGEPAL, CA-630, in water). Resulting nuclei were centrifuged at 500 rcf for 10 minutes. Pelleted nuclei were resuspended in 50 µl transposase reaction mix (1x Tagment DNA buffer (Illumina 15027866), 2.5 µl Tagment DNA enzyme I (Illumina 15027865), and incubated at 37°C for 30 min on a heat block. For isolations resulting in under 30,000 microglia, microglia directly placed in 50 µl transposase reaction mix, as indicated above and incubated for 37°C for 30 min. DNA was purified with Zymo ChIP DNA concentrator columns (Zymo Research D5205), eluted with 11 µl of elution buffer, and amplified using NEBNext High-Fidelity 2x PCR MasterMix (New England BioLabs M0541) with the Nextera primer Ad1 (1.25 µM) and a unique Ad2.n barcoding primer (1.25 µM) for 8-12 cycles. Libraries were size-selected through gel excision for fragments that were 175-255 bp and single-end sequenced for 51 cycles on a HiSeq 4000 or NextSeq 500.

### Chromatin immunoprecipitation-sequencing (ChIP-seq)

FACS-purified microglia were spun down at 300 rcf and resuspended in 1% PFA. After rocking for 10 minutes at room temperature, PFA was quenched with 1:20 2.625M glycine for 10 minutes at room temperature. Fixed cells were washed twice and centrifuged at 800-1000 rcf for 5 minutes and snap frozen in liquid nitrogen. Snap-frozen microglia (750,000) were thawed on ice, resuspended in 130 µl of LB3 buffer (10 mM TrisHCl pH 7.5, 100 mM NaCl, 1 mM EDTA, 0.5 mM EGTA, 0.1% Na-Deoxycholate, 0.5% N-Lauroylsarcosine, 1x protease inhibitors), and transferred to microtubes with an AFA Fiber (Covaris, MA). Samples were sonicated using a Covaris E220 focused-ultrasonicator (Covaris, MA) for 12 cycles of 60 secs (Duty: 5, PIP: 140, Cycles: 200, AMP/Vel/Dwell: 0.0). The sonicated sample was transferred to an Eppendorf tube, to which Triton X-100 was added to achieve a final concentration of 1%. Supernatant was spun at 21,000 rcf and the pellet discarded. 1% of the total volume was saved as DNA input control and stored at -20°C until library preparation. For the immunoprecipitation, 25 µl of Protein A DynaBeads (Thermo Fisher Scientific 10002D) and 1 µl of H3K27ac antibody (Active Motif, 39085) were added to the supernatant and rotated at 4°C overnight. Dynabeads were washed 3 times with Wash Buffer 1 (20 mM Tris-HCl pH 7.4, 150 mM NaCl, 2 mM EDTA, 0.1% SDS, 1% Triton X-100), three times with Wash Buffer 3 (10 mM Tris-HCl pH, 250 mM LiCl, 1 mM EDTA, 1% Triton X100, 0.7% Na-Deoxycholate), three times with TET (10 mM Tris-HCl pH 8, 1 mM EDTA, 0.2% Tween20), once with TE-NaCl (10 mM Tris-HCl pH 8, 1 mM EDTA, 50 mM NaCl) and resuspended in 25 µl TT (10 mM Tris-HCl pH 8, 0.05% Tween20). Input samples were adjusted to 25 µl with TT. Sample and input libraries were prepared using NEBNext Ultra II DNA Library Prep kit (New England BioLabs E7645) according to manufacturer’s instructions. Samples and inputs were de-crosslinked (RNase A, Proteinase K, and 4.5 µl of 5M NaCl) and incubated overnight at 65°C. Libraries were PCR-amplified using NEBNext High Fidelity 2X PCR MasterMix (New England BioLabs M0541) for 14 cycles. Libraries were size-selected through gel excision for fragments that were 225 to 500 bp and single-end sequenced for 51 cycles on a HiSeq 4000 or NextSeq 500.

### mRNA isolation

Snap-frozen human fetal and postnatal cortical tissue were placed in TRIzol LS (Life Technologies) and homogenized using Powergen 125 homogenizer (Thermo Scientific). FACS-sorted cells were stored in TRIzol LS. Total RNA was extracted from homogenates and cells using the Direct-zol RNA MicroPrep Kit (Zymo Research R2052) and stored at - 80°C until RNA-seq cDNA libraries preparation.

### RNA-seq library preparation

RNA-seq libraries were prepared as previously described^9^. Briefly, mRNAs were incubated with Oligo d(T) Magnetic Beads (New England BioLabs S1419), then fragmented in 2x Superscript III first-strand buffer (Thermo Fisher Scientific with 10mM DTT (Thermo Fisher Scientific 18080044) at 94°C for 9 minutes. Fragment mRNA was then incubated with 0.5 μl of Random primers (3 mg/mL) (Thermo Fisher Scientific 48190011), 0.5 μl of 50mM Oligo dT primer, (Thermo Fisher Scientific, 18418020), 0.5 μl of SUPERase-In (Thermo Fisher Scientific AM2694), 1 μl of dNTPs (10 mM) at 50°C for one minute. Then, 1 μl of 10mM DTT, 6 μl of H_2_O+0.02%Tween-20 (Sigma), 0.1 μl Actinomycin D (2 mg/mL), and 0.5 μl of Superscript III (Thermo Fisher Scientific) were added to the mixture. cDNA was then generated by incubating mixture in a PCR machine at the following conditions: 25°C for 10 minutes, 50°C for 50 minutes, and a 4°C hold. Product was purified using RNAClean XP beads (Beckman Coulter A63987) according to manufacturer’s instructions and eluted with 10 μl of nuclease-free H_2_O. Eluate was then incubated with 1.5 μl of Blue Buffer (Enzymatics P7050L), 1.1 μl of dUTP mix (10 mM dATP, dCTP, dGTP and 20 mM dUTP), 0.2 mL of RNase H (5 U/mL Y9220L), 1.2 μl of H_2_O+0.02%Tween-20, and 1 μl of DNA polymerase I (Enzymatics P7050L) at 16°C overnight. DNA was then purified using 3 μl of SpeedBeads (Thermo Scientific Fisher 651520505025) resuspended in 28 μl of 20% PEG8000/2.5M NaCl to final of 13% PEG. DNA eluted with 40 mL nuclease free H_2_O+0.02%Tween-20 and underwent end repair by blunting, A-tailing and adaptor ligation as previously described^10^ using barcoded adapters. Libraries were PCR-amplified for 12-15 cycles, size selected by gel extraction for 200-500 bp, and sequenced on a HiSeq 4000 (Illumina) or a NextSeq 500 (Illumina) for 51 cycles.

### scRNA-seq data generation

Sorted microglia were centrifuged for 5 minutes at 300 rcf and the supernatant was carefully aspirated leaving approximately 25 μl behind. Cells were resuspended in up to 40 μl reaction buffer (0.1% BSA (SRE0036-250ML, Sigma) and 1 U/μl RNasin inhibitor (PAN21110, Promega) in PBS (21-040-CV, Corning)). An aliquot of the cell suspension was mixed with Trypan Blue (T10282, Invitrogen) to count and check viability using a hemocytometer. 12,000 cells (viability 65-100%) were loaded onto a Chromium Controller (10x Genomics). Libraries were generated according to manufacturer specifications (Chromium Single Cell 3’ Library and Gel Bead Kit v3, 1000075; Chromium Single Cell 3’ Library Construction Kit v3, 100078; Chromium Chip B Single Cell Kit, 1000153; Single Index Kit T Set A, 1000213). cDNA was amplified for 12 PCR cycles. SPRISelect reagent (Beckman Coulter) was used for size selection and clean-up steps. Final library concentration was assessed by Qubit dsDNA HS Assay Kit (Thermo-Fischer Scientific) and fragment size was checked using the High Sensitivity D1000 ScreenTape assay on a Tapestation 4200 (Agilent) to ensure that fragment sizes were distributed normally about 500 bp. Libraries were sequenced using a NextSeq 500 or NovaSeq 6000 (Illumina) using these read lengths: Read 1: 28 cycles, Read 2: 91 cycles, Index 1: 8 cycles.

### scATAC-seq data generation

Sorted microglia were centrifuged for 5 minutes at 300 rcf and the supernatant was carefully aspirated. Cells were permeabilized using lysis buffer (10 mM Tris-HCl pH 7.4 (15567027, Thermo Fischer Scientific), 10 mM NaCl (ICN15194401, Fischer Scientific), 3 mM MgCl_2_ (194698, Mp Biomedicals Inc.), 0.1% Tween 20 (P7949, Sigma), 0.1% IGEPAL-CA630 (I8896, Sigma), 0.01% Digitonin (G9441, Promega), and 1% BSA in nuclease free water) and incubated for 5 minutes on ice. Permeabilized nuclei were washed using washing buffer (lysis buffer without IGEPAL-CA630 and Digitonin), centrifuged for 5 minutes at 500 rcf and resuspended in Nuclei buffer (10x Genomics). An aliquot was mixed with Trypan Blue and counted using a hemocytometer. Up to 15,300 nuclei were tagmented before loading onto a Chromium Controller; libraries were generated according to manufacturer specifications ((Chromium Next GEM Single Cell ATAC Library and Gel Bead Kit v1.1, 1000175; Chromium Next GEM Chip H Single Cell Kit, 1000162; Single Index Kit N Set A, 1000212, 10x Genomics)). Libraries were amplified for 10 PCR cycles. SPRISelect reagent (Beckman Coulter B23318) was used for size selection and clean-up steps. Final library concentration was assessed by Qubit dsDNA HS Assay Kit (Thermo-Fischer Scientific) and fragment size was inspected using the High Sensitivity D1000 ScreenTape assay on a Tapestation 4200 (Agilent). Libraries were sequenced using a NextSeq 500 or NovaSeq 6000 (Illumina) using these read lengths: Read 1: 50 cycles, Read 2: 50 cycles, Index 1: 8 cycles, Index 2: 16 cycles.

## Data analysis

### Data preprocessing

Raw reads were obtained from Illumina Studio pipeline. ATAC libraries were trimmed to 30bp. RNA-seq data was mapped to hg38/mm10 genome using STAR^11^ with default parameters. H3K27acetylation ChIP-seq and ATAC-seq data were mapped to hg38 genome using bowtie2^12^ with default parameters. Finally, HOMER^10^ tag directories were created for mapped samples.

### External RNA-seq data

Data from Matcovitch-Natan *et al*.^13^, Bian et al^14^, Hasselmann et al.,^15^, and Svoboda et al.^16^ were obtained via the sequence read archive (SRA)^17^ and were preprocessed with the same pipeline as described above.

### RNA-seq data analysis

HOMER’s analyzeRepeats.pl was used to calculate gene expression raw counts with the option “-condenseGenes -count exons -noadj” and transcript per kilobase million (TPM) with the option “-count exons -tpm”. Genes shorter than 200bp were removed. TPMs were matched to raw counts using accession numbers. Differentially expressed genes comparisons using human microglia, human cortical, iMGs, oMGs data were assessed with DESeq2^18^ at an adjusted p-value < 0.05 and fold-change > 2 where indicated. Multiple testing correction was adjusted using the procedure of Benjamini and Hochberg under DESeq2 framework.

Mapping of orthologous genes between mouse and human was done using the one-to-one orthologs from the Ensembl (version 84) Compara database.

### Aggregated expression score

The expressions of mouse orthologs that are associated with human microglia differentially expressed genes were used to compute aggregated expression score (AES). For each gene set and each mouse developmental time point, AES is calculated as the mean expression of genes that are in the enriched GO terms divided by the mean expression of all genes. Random gene set containing 750 random genes was used as a control and its AES is computed together with enriched GO terms.

### NicheNet Analysis

NicheNet models the influence of ligands expressed by a sender cell on a set of target genes in a receiver cell using a model of intracellular signaling linking receptors to target genes (available at https://github.com/saeyslab/nichenetr)^19^. NicheNet assesses whether a given ligand could drive transcription of a set of target genes relative to all expressed genes within a cell or tissue. In this study, we used NicheNet to identify ligands expressed by microglia (autocrine) and bulk cortex tissue (paracrine) that could drive fetal-specific or postnatal specific microglial gene expression. Genes that were differentially expressed at an p-adj. < 0.05 and fold change > 2 between conditions were used to establish the target set of genes for indicated states compared (e.g. the fetal and postnatal state), with all other genes serving as the background set. The NicheNet ligand-target model was filtered to only include ligands and receptors expressed at a level of >5 TPM for both autocrine and paracrine analysis. Additionally, the NicheNet model was filtered to only include ligand-receptor interactions annotated by curated databases and exclude ligand-receptor interactions annotated by protein-protein interaction databases. Ligands that were expressed at least 8-fold higher in microglia compared to the corresponding developmental age-matched cortex were considered as likely autocrine signals, whereas ligands expressed at less than a than 2-fold increase in respective cortex versus microglia comparisons were considered to act on microglia in both an autocrine and paracrine fashion. Assignment of upstream receptors to downstream ligands in Figure 2A was performed manually using prior literature to support connections.

### Weighted Correlation Network Analysis

Weighted Correlation Network Analysis (WGCNA) was performed using TPMs calculated using defined by HOMER’s analyzeRepeats.pl function. Samples’ tissue source (microglia or whole cortex), developmental age (fetal or postnatal) and gender were used as input traits. Soft threshold was set to 10 to reach 0.90 of scale free topology index when constructing the gene co-expression matrix. Minimum module size was set to 100 and maximum dissimilarity was set to 0.01 when merging modules. Module-trait correlations were computed using Pearson’s correlation, modules with correlation coefficients p>0.7 or p<-0.7 and p-value <0.05, for at least one trait, were considered as significant and subjected to functional annotation. Gender trait was not used in further analysis as no gene module is significantly correlated with gender. Next, we computed the Kleinberg’s hub centrality scores for genes in each significant module. We performed Gene Ontology (GO) enrichment analysis on genes in the significant modules for functional annotation. Bioconductor package *topGO* was used to identify significantly enriched GO terms associated with genes in each module, the p-value and q-value cutoffs were set to 0.01 and 0.05, respectively. The enriched GO terms were used to annotate each gene module.

### ATAC-seq analysis

Peaks were called using HOMER’s findPeaks command with the following parameters: “-style factor -size 200 - minDist 200”. Peaks were merged with HOMER’s mergePeaks and annotated using HOMER’s annotatePeaks.pl using all tag directories. Subsequently, DESeq2^18^ was used to identify the differentially chromatin accessible distal sites (3000bp away from known TSS) or proximal sites (<500bp away from known transcript) with p-adj < 0.05 and fold change > 2.

### ChIP-seq analysis

ChIP-seq peaks were called using “findPeak” command from HOMER with the following parameters “-style histone -size 200 -L 0 -C 0 -fdr 0.9”.

### Motif analysis

*De novo* motif analysis was performed using HOMER’s findMotifsGenome.pl with random genome sequences as background peaks. Motif enrichment scoring was performed using binomial distribution under HOMER’s framework.

### Single cell RNA-seq processing

Raw sequencing data was demultiplexed and preprocessed using the Cell Ranger software package v3.0.2 (10x Genomics). Raw sequencing files were first converted from Illumina BCL files to FASTQ files using cellranger mkfastq for scRNA-seq. Demultiplexed FASTQs were aligned to the GRCh38 reference genome (10x Genomics),and reads for exonic reads mapping to protein coding genes, long non-coding RNA, antisense RNA, and pseudogenes were used to generate a counts matrix using Cellranger count; expect-cells parameter was set to 5,000. Fastq files were processed using Cell Ranger 3.0.2 software to process barcodes and demultiplex reads using default parameters. The resulting count matrices were filtered and analyzed using Seurat package (version 3.2.2). Genes expressed in fewer than 3 cells and cells expressing fewer than 500 genes were removed from further analysis. Normalization was performed using the Seurat default parameters. We used Harmony (https://github.com/immunogenomics/harmony)^20^ to integrate samples. Louvain algorithm with resolution=0.3 from Seurat package was applied to cluster cells. The result was projected on t-distributed stochastic neighbor embedding (tSNE) for visualization.

### RNA velocity analysis

RNA velocity analysis was performed using the Velocyto package^21^ (available at velocyto.org). Fastq files were trimmed and mapped using STAR to hg38 reference genome. The intronic and exonic information was extracted; and the ratios of the spliced and unspliced variants of each gene in every cell were computed; changes in these ratios for a gene in a cell allow for inferences in cell state changes among the population. Arrows point to the position of the future state and to infer cell trajectories.

### Enhancer associated transcription factor activity (Extended Data Figure 8a)

#### Enhancer calling

Active enhancers were selected using the following criteria: 1) open chromatin region marked by ATAC-seq peak, 2) located outside of transcription start site (−2kb to +100bp), and 3) have at least 16 tag counts in the H3K27ac peak overlapping with the ATAC-seq peak. Peak calling and annotation were performed using Homer findPeaks and annotatePeaks.pl functions, as previously described. Fetal and postnatal specific enhancers are defined as enhancers that are not overlapped between fetal and postnatal groups.

#### Transcription Factor Motif

Instances of transcription factor motifs in enhancers were defined using Homer findMotifsGenome.pl function with -find flag. The reference motif library for motif identification was downloaded from Jaspar non-redundant vertebrates motif library (http://jaspar.genereg.net/downloads/) and converted using MAGGIE^22^. We used the distance method to assign enhancers to target genes. Motifs identified in enhancers that were assigned target genes were then used to construct transcription factor – target gene networks (TF-gene networks). Only transcription factors with mean expression at least 16 TPM from either bulk fetal RNA-seq or postnatal RNA-seq dataset were considered. Separate TF-gene networks were constructed for fetal and postnatal groups.

#### TF module activity score calculation

We used the fetal and postnatal microglia TF-gene networks and scRNA-seq expression data, normalized using Seurat, to compute activity score for each TF for each developmental age group. Activity score for each TF is computed using VISION^23^ calcSignatureScores function with TF -gene network and scRNA-seq expression data as inputs.

### Single-cell ATAC-Seq preprocessing and clustering

Raw sequencing files were first converted from Illumina BCL files to FASTQ files using cellranger-atac mkfastq. Demultiplexed FASTQs were aligned to the GRCh38 reference genome (10x Genomics) to identify chromatin accessible peaks using cellranger-atac count. Sequencing reads of the four donors were demultiplexed and processed using the Cell Ranger software package Cellranger-atac v1.1.0 (10x Genomics). Reads were aligned to the human reference hg38 (Cell Ranger software package cellranger-atac-1.1.0/bwa/v0.7.17). The fragment files generated by Cell Ranger were then tagged by read and donor and combined into a unique fragment file. We then computed a Transcription Start Site (TSS) enrichment score for each cell using +/- 2kbp from the TSS as reference promoter regions. We used a flank size of 100pb at the beginning and end of the promoters, a smoothing window of 10bp, and a TSS region of 50bp to infer the maximum TSS enrichment for each cell. We called peaks on the merged fragment file using MACS2^24^ with the following options: -nomodel -keepdup-all -q 0.01 --shift 37 --extsize 73. We then filtered the top 5% of the peaks, merged the overlapping peaks, and transformed each peak using its center with +/1000bp as boundaries. We filtered cells with less than 1500 fragments and with TSS < 7 and created a binary sparse *cell x peaks* matrix. We converted the sparse matrix into a lower dimensional embedding using a Latent Semantic Analysis (LSA) approach by weighting the features with a tf-idf scheme and extracted 25 new dimensions using a Singular Value Decomposition (SVD) algorithm using RobustSVD and TfIdfTransformer classes from the scikit-learn package^25^. We used the new embedding and the batch ID of each cell as input for Harmony^20^ and inferred a new embedding (ncells *x 25*) corrected for the batch effect from the donor ID. We then projected the new embeddings with and without harmony correction into a 2D space using UMAP (umap-learn package) using correlation as similarity, 2.0 as repulsion strength, and 0.01 as min distance^26^. We clustered the UMAP spaces using HDBSCAN algorithm (python hdbscan package https://hdbscan.readthedocs.io/en/latest/). We called peaks on each individual cluster obtained by this procedure by agglomerating the reads of the cells according to their labels and used MACS2 with the setting described above. We merged using bedtools the peak list of each cluster into a final set of 129576 peaks. We performed the same workflow described above with the 129576 peaks as input features to create a cells x features binary matrix, convert into a lower dimensional embedding, and correct batch effects with Harmony. We constructed a K-Nearest Neighbor (KNN) similarity graph with K=50 and similarity=cosine using the scikit-learn NearestNeighbors class. Finally, we clustered the similarity graph with the Louvain clustering algorithm^27^ with different resolutions R (1.0, 1.5, and 2.0) from the python-louvain package (https://github.com/taynaud/python-louvain), and selected R=1.5 (6 clusters) based on the sum(-log10(fisher p-values)) of the significant cluster features.

### Single-cell motif enrichment analysis

We used ChromVAR (https://greenleaflab.github.io/chromVAR/)^28^ to compute motif enrichment at the single-cell level. We used the merged list of cluster peaks (center +-250pb), and a list of 870 of non-redundant reference motifs as input for chromVAR workflow. We identified differentially enriched motifs for each cluster using the following strategy: for each cluster and each motif, we computed a Rank Sum test between the ChromVAR Z-score distributions within and outside the cluster. We applied a Benjamini-Hochberg FDR correction on the p-values.

### Multi-modal correlation matrices

Using the 6 clusters C obtained with R=1.5 and the 129576 peaks P we transformed the cells x peaks sparse binary matrix into a clusters x peaks matrix by taking the average number of cells having a peak p accessible within a cluster c for each p from P and c from C. Each column and row are then scaled to have a norm equal to 1. Using the same strategy, we created a donors x peaks using the donor ID (four donors) of each cell as label. Finally, we created a bulks x peaks matrix by counting the number of reads overlapping each peak p from P for the 30 bulk ATAC-Seq datasets. We then scaled the matrices (each column has mean=0 and std=1) and computed the clusters x bulk and donors x bulk kernel similarity matrices using the Pearson correlation as metric. Finally, we plotted the similarity matrices using the clustermap function from the seaborn python library (https://seaborn.pydata.org/api.html).

## Statistical analyses

Gene expression differences were calculated with DESeq2 with Benjamini-Hochberg multiple testing correction. Genes are considered differentially expressed at >2FC, p.adj < 0.05.

## Data Visualization

Heatmaps were generated with the pheatmap packages in R and other plots were made with ggplots2 in R with colors reflecting the scores/expression values, including z-scores, as noted in each figure. Circo plots showing genes related to neurodevelopmental diseases and for NicheNet ligand-receptor analysis were generated using the R package Circlize^29^ package in R. Violin plots for scRNA-seq data was produced using Vlnplot function from Seurat package. Bar charts were generated using Prism 7.0 and presented as mean ± SEM. Browser images were generated from the UCSC Genome Browser.

## Data Availability

Previously reported data were downloaded from GEO: Matcovitch-Natan *et al*.^13^ (GSE79812), Bian et al^14^ (GSE110611), Hasselmann et al.^15^ (GSE133432), and Svoboda et al.^16^ (GSE139192). A portion of the human postnatal microglia data are available through NCBI dbGaP (phs001373.v1.p1).

## Code availability

Code for WGCNA, aggregated expression score analysis, scRNA-seq velocity analysis, enhancer associated transcription activity, and TROM analysis are available here: https://github.com/rzzli/FetalMicroglia. Code for scATAC-seq BAM/BED files processing, sparse matrix creation, TSS enrichment computation, matrix clustering and visualization are available here: https://gitlab.com/Grouumf/ATACdemultiplex. Description of the custom set of the 870 non-redundant motifs used as input for chromVAR analysis is described here: https://github.com/GreenleafLab/chromVARmotifs.

## Acknowledgements

We thank the UC San Diego Center for Perinatal Discovery for providing fetal brain samples for this study. We thank L. Van Ael for assistance with manuscript preparation. We thank Uri Manor for imaging assistance. C.Z.H. is supported by the Cancer Research Institute Irvington Postdoctoral Fellowship Program. N.G.C. is supported by NIH K08 NS109200, The Hartwell Foundation, and the Doris Duke Foundation. This work was supported in part by the Flow Cytometry Core Facility of the Salk Institute with funding from NIH-NCI CCSG: P30 014195 and Shared Instrumentation Grant S10-OD023689 (Aria Fusion cell sorter). Work at the Center for Epigenomics was supported in part by the UC San Diego School of Medicine. This publication includes data generated at the UC San Diego IGM Genomics Center utilizing an Illumina NovaSeq 6000 that was purchased with funding from a National Institutes of Health SIG grant (#S10 OD026929)

## Author Contributions

C.Z.H., R.Z.L, C.K.G. and N.G.C. conceived the project and wrote the manuscript with input from co-authors. N.G.C., L.L., M.G., S.B., D.D.G, and M.L.L. were involved in patient identification, consent and tissue acquisition. C.Z.H. and N.G.C. isolated cells which was sorted by C.C. Sequencing libraries were prepared by C.Z.H. and J.B and analyzed by C.Z.H., R.Z.L., H.B., O.P., J.B., J.C., S.B., A.W. B.R.F. provided experimental assistance. E.H. performed confocal imaging. E.H., A.S.W., S.S., R.K., C.N., D.S., G.R., S.A. and A.J. performed in vitro experiments and immunohistochemistry. E.S. provided adult mouse cortex data. The project was supervised by C.K.G. and N.G.C. All authors contributed to editing and review of the manuscript.

## Supplementary information

Supplementary information is available for this paper.

**Supplementary Figure 1**. FACS gating strategy for sorting oMGs and iMGs.

**Supplementary Table 1:** Clinical information of postnatal microglial samples. Table includes brain region from which microglia were isolated, patient age/gender, the clinical pathological diagnosis of resected tissue, antiepileptics patients were receiving at the time of surgery, and whether stereoelectroencephalography was performed with implantable electrodes relative to surgical dates.

**Supplementary Table 2**: Full list of gene modules clustered by WGCNA. Annotation represents the GO analysis based on genes within the category.

**Supplementary Table 3:** Signature scores correlating fetal and postnatal microglial human GO categories to murine microglia RNA-seq across developmental timepoints.

## Extended Data Figure Legends

**Extended Data Figure 1. Transcriptomes of human fetal and postnatal microglia and brain tissue**.

a. Flow cytometry panel depicting sorting strategy for human fetal microglia. After live-dead and singlets gating, fetal microglia are defined by CD11b+CD45+CX3CR1+CD64+.

b. Immunohistochemistry of fetal brain illustrating colocalization of PU.1 (white) with IBA1 (red) positive cells and neurons, indicated by MAP2 staining (green). Microglia were identified throughout all brain regions.

c. Bar charts depicting the number of samples per gestational week processed and analyzed for the indicated assays. Numbers for microglia are depicted in the left and middle panel while bulk fetal cortex is represented in the right panel.

d. Metascape enrichment analysis of differentially expressed genes between human fetal and postnatal microglia. Top enriched gene modules are shown, x-axis is the -log10(p) of the enrichment levels.

**Extended Data Figure 2. Monogenic neurodevelopment disorders gene expression in fetal microglia**.

a. Pie charts depicting the number of monogenic NDD genes in indicated NDDs that are significantly expressed in fetal microglia (blue), fetal cortex (green), or similarly expressed (grey).

b. Heatmaps of the expression levels of monogenic NDD genes for indicated NDDs. Black side bar denotes genes that are significant (> 2FC, p-adj < 0.05) while grey indicates no statistical significance.

**Extended Data Figure 3. Expression differences of monogenic neurodevelopment disorder gene expression between mouse and human**.

a. Bar chart of TPM expression levels of select genes unique to either fetal or postnatal microglia enriched genes.

b. Bar charts of TPM expression levels of cytokines essential for microglia survival and function.

c. Immunohistochemistry of fetal brain sections for IL17R in microglia and IL17 expression in fetal cortex adjacent to microglia.

d. Circos plot of monogenic genes associated with indicated disorders differentially expressed in human fetal (blue) compared to postnatal microglia (red) (outer most track). Each inner track shows the status of these genes’ ortholog in mouse differential expression analysis, where blue is higher expressed in the mouse microglia of the specified age compared to adult mouse microglia (red). Mouse ages progress as follows (outer to inner: E10.5, E12.5, E14, E16.5, PN3, PN6, and PN9). Grey denotes gene expression is nonsignificant between the indicated mouse microglia age and adult mouse microglia.

*adj. p-val<0.05, ** adj. p-val <0.01, *** adj. p-val <0.001, **** adj. p-val <0.0001

**Extended Data Figure 4. Select markers of chromatin accessibility in scATAC-seq of human microglia**.

a. Heatmap depicting correlation of peaks generated from pseudobulk of scATAC-seq samples with bulk ATAC-seq samples.

b. Activities of TFE3, EGFR3, MAFB and KLF6 motifs measured by ChromVar and projected onto UMAP space.

**Extended Data Figure 5. Cluster analysis of scRNA-seq of human fetal and postnatal microglia I**.

a. RNA velocity analysis on scRNA-seq of fetal and adult microglia (left) and tSNE projection of relative expression for indicated genes (right).

b. tSNE plots of expression of canonical microglia genes.

c. Violin plots depicting log transformed expression level of ligand-receptor pairs, IGF1-IGF1R, CSF1-CSF1R, across clusters.

d. Expression of genes enriched in the immune modulatory cluster (cluster 1,top; cluster 3, bottom) from scRNA-seq data of human fetal and postnatal microglia, as projected onto tSNE plots or as log transformed values in violin plots across clusters.

**Extended Data Figure 6. Cluster analysis of scRNA-seq of human fetal and postnatal microglia II**.

a. Bar charts showing TPM expressing levels of cell cycle genes implicated in intellectual disability.

b. Immunohistochemistry of fetal microglia for KI67 as a marker of cell division.

c. tSNE plots of expression of genes in S6A, showing that many of them are enriched in the cell cycle cluster.

d. tSNE plots of relative expression of genes enriched in interferon responsive group (cluster 6) (left) and violin plots of log transformed expression level of select genes (right).

e. tSNE plots of relative expression of genes associated with neural progenitor cells, suggestive of a phagocytic group. Inset of tSNE plot of *CSF1R* expression shows that this phagocytic group also co-express CSF1R, a microglia marker.

f. Immunohistochemistry of IBA1+ (green) fetal microglia engulfing a NPC, marked by SOX2 (red) staining.

*adj. p-val<0.05, ** adj. p-val <0.01, *** adj. p-val <0.001, **** adj. p-val <0.0001

**Extended Data Figure 7. Detection of human border-associated brain macrophages and monocytes in scRNA-seq**.

a. tSNE plots of relative expression of genes enriched in border-associated macrophage (BAM) group (cluster 5, left) and monocytes (cluster 8, right). Violin plot depicting log transformed expression level of select cluster marker genes across clusters (middle).

b. Immunohistochemistry of fetal brain tissue showing colocalization of LYVE1 positive cells with tissue border, detected by Laminin staining.

c. Immunohistochemistry of fetal cortical tissue (top) and brainstem (bottom) for border-associated macrophages, using LYVE1 and IBA1 (top) and CD163 and IBA1 (bottom).

**Extended Data Figure 8. Strategy for enhancer associated transcription factor activity and expanded by cluster TF activity analysis**.

a. Schematic of workflow to compute TF activity scores.

b. Heatmap of TF activity scores of scRNA-seq data grouped by cluster.

c. TF activity scores for SPIB, FOSL1::JUNB, MAFB and IRF5, projected onto tSNE space.

**Extended Data Figure 9. Differential gene expression and chromatin accessibility between oMGs and iMGs**.

a. MA plot of gene expression between iMG and oMG. Genes that are significantly differential (> 2-fold, FDR < 0.05) are highlighted in green (iMG) and purple (oMG).

b. Metascape enrichment analysis of differentially expressed genes between iMG and oMG. Top enriched gene modules are shown, x-axis is the -log10(p) of the enrichment levels.

c. Volcano plot of distal (>3kb from TSS) ATAC-seq peaks between iMG and oMG. Colored points indicate significantly differential peaks (> 2-fold, FDR < 0.05) with those enriched in iMG in green and those enriched in oMGs in dark blue (left). Motif enrichment analysis of distal differential accessible chromatin regions enriched in iMGs shown, using genomic sequence as background (right).

**Extended Data Figure 10. Expression of genes related to human microglia function and NDDs in iMGs and oMGs**.

a. Bar charts of TPM expression levels of genes associated with lysosomal storage disease in model system (oMG, iMG, and bulk organoids, top panel) and primary tissue (human fetal and postnatal microglia, fetal and postnatal bulk cortex, bottom panel).

b. Bar charts of TPM expression levels of NPC genes in primary tissue (human fetal and postnatal microglia, fetal and postnatal bulk cortex) and model system (oMG, iMG, and bulk organoids).

c. Bar charts of TPM expression levels of *IL34, CSF1*, and *CSF1R* in primary tissue (human fetal and postnatal microglia, fetal and postnatal bulk cortex) and model system (oMG, iMG, and bulk organoids) (left). UCSC genome browser tracks (right) of RNA-seq, ATAC-seq, and H3K27acetylation of *CSF1R* in human fetal and postnatal microglia and RNA-seq of *CSF1R* in iMGs and oMGs.

*adj. p-val<0.05, ** adj. p-val <0.01, *** adj. p-val <0.001, **** adj. p-val <0.0001

