## Supplemental Files for "Gene regulatory networks underlying human microglia maturation"

#### Extended Data Figure 1

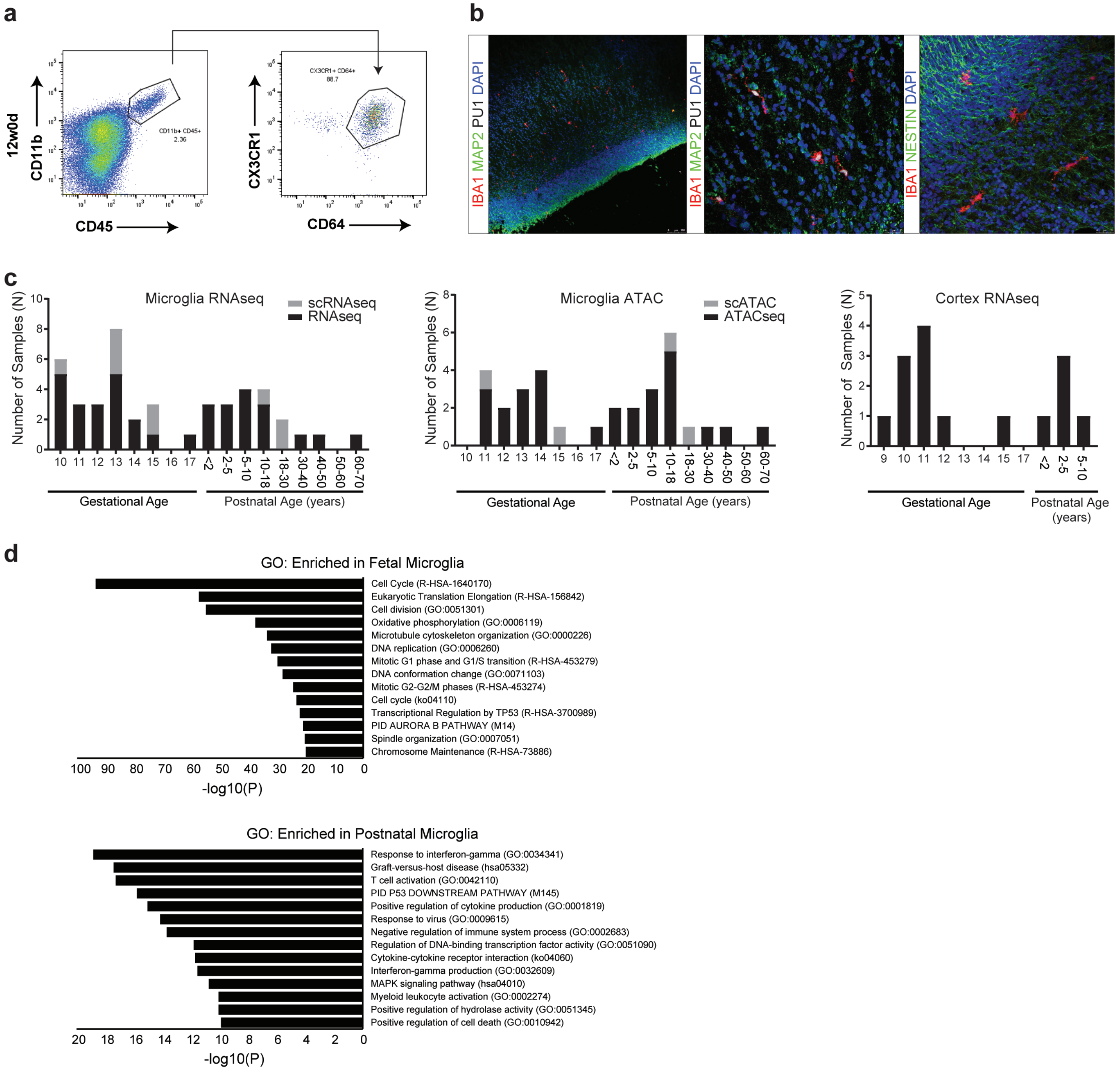

### Extended Data Figure 2

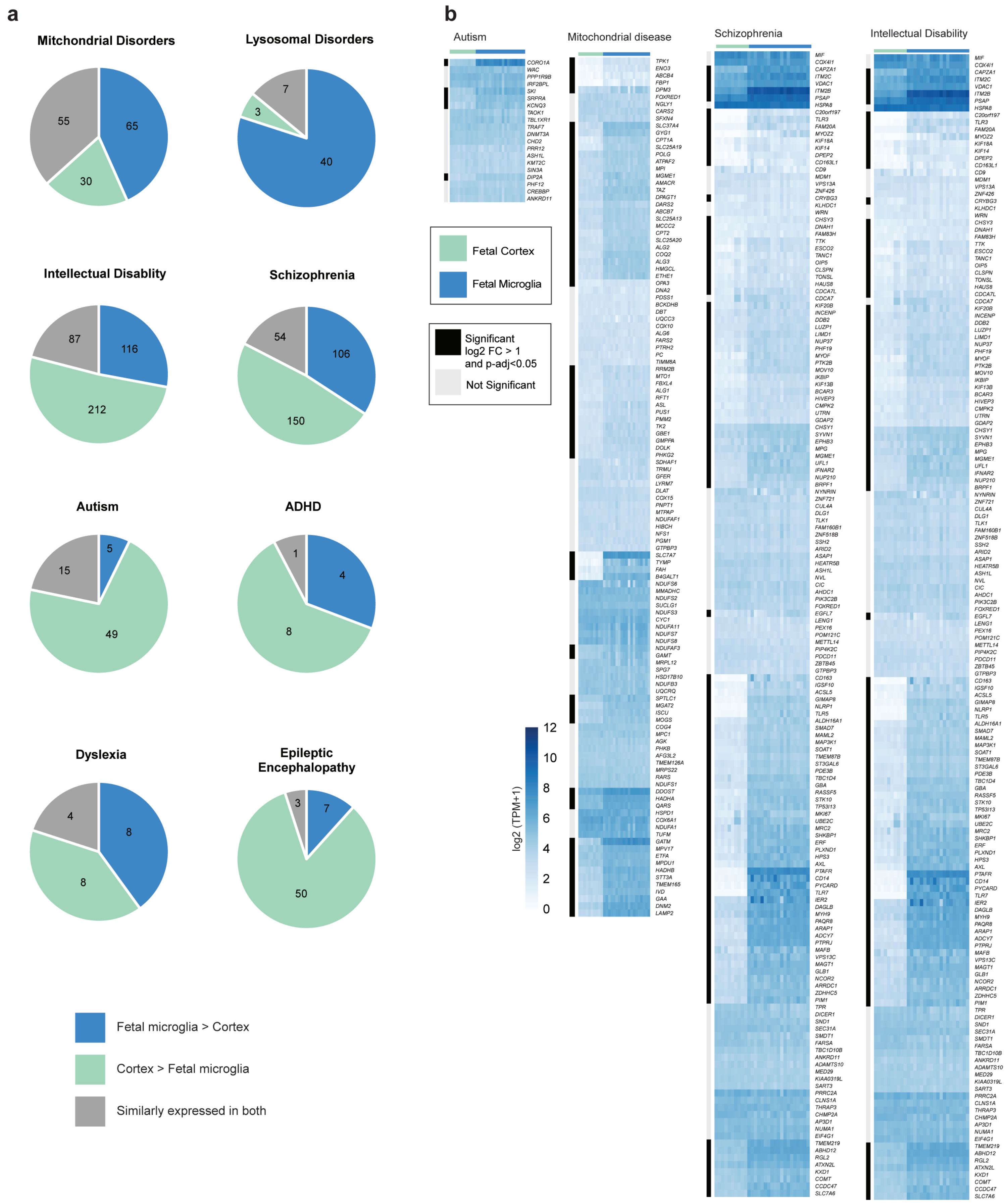

**Extended Data Figure 3**

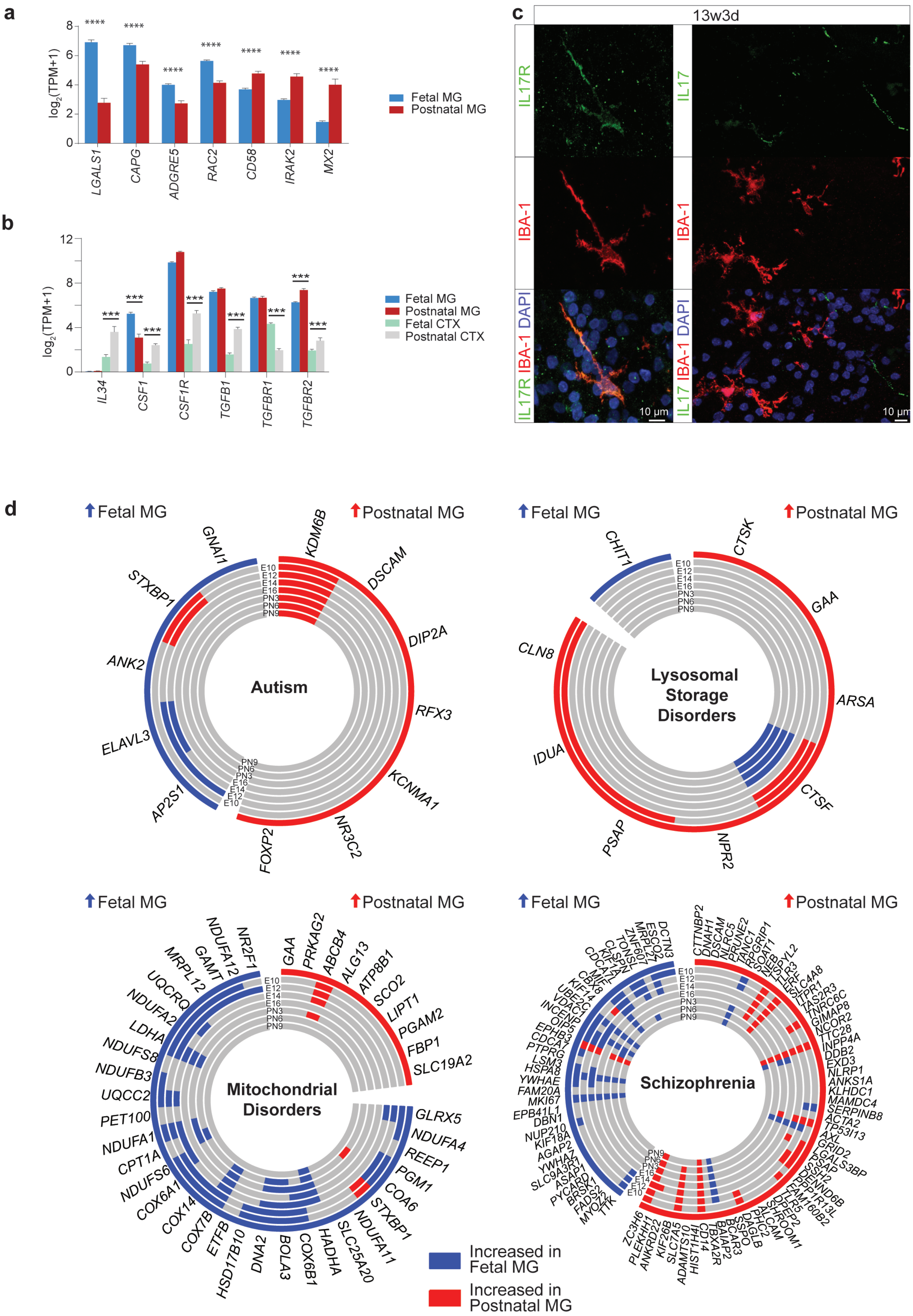

### Extended Data Figure 4

**a**

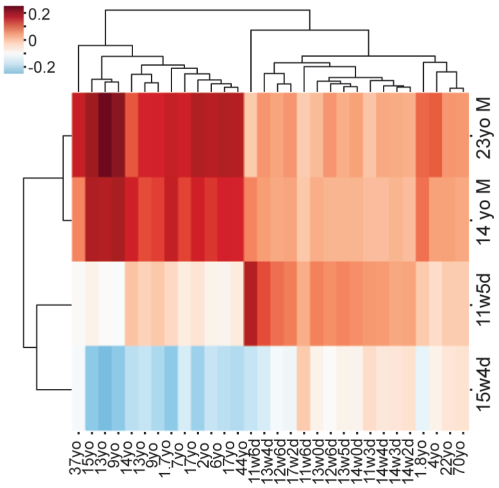


**b**

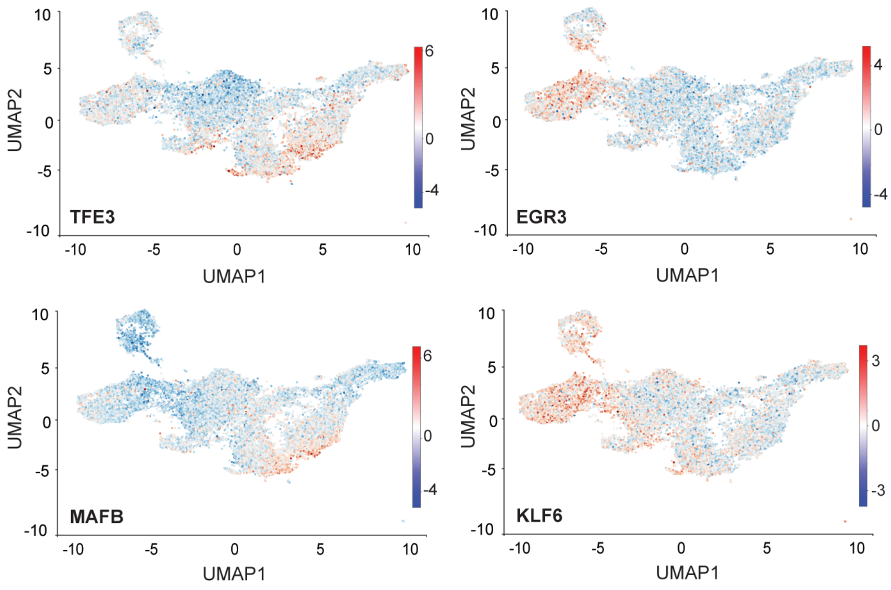

#### Extended Data Figure 5

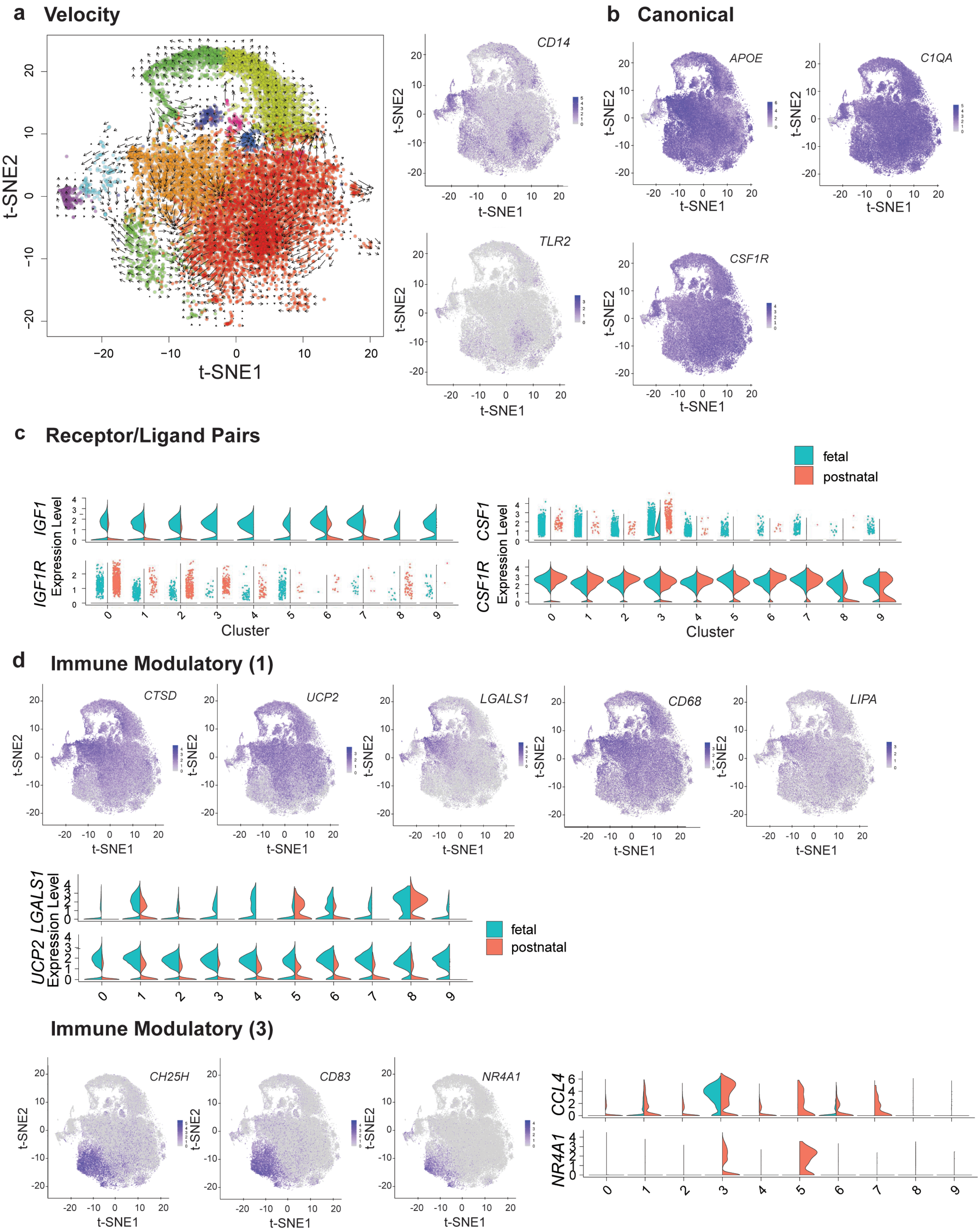

#### Extended Data Figure 6

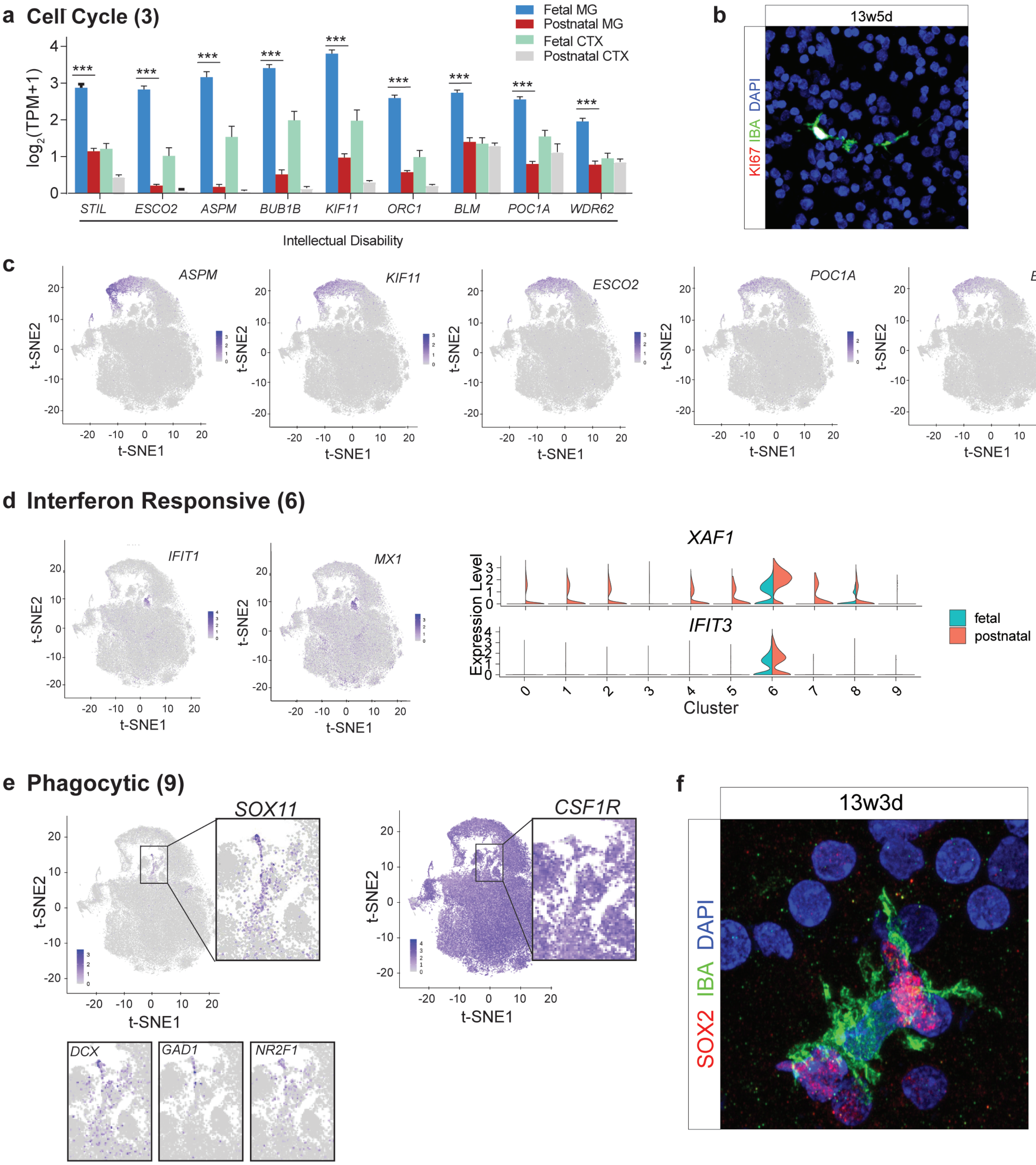

### Extended Data Figure 7

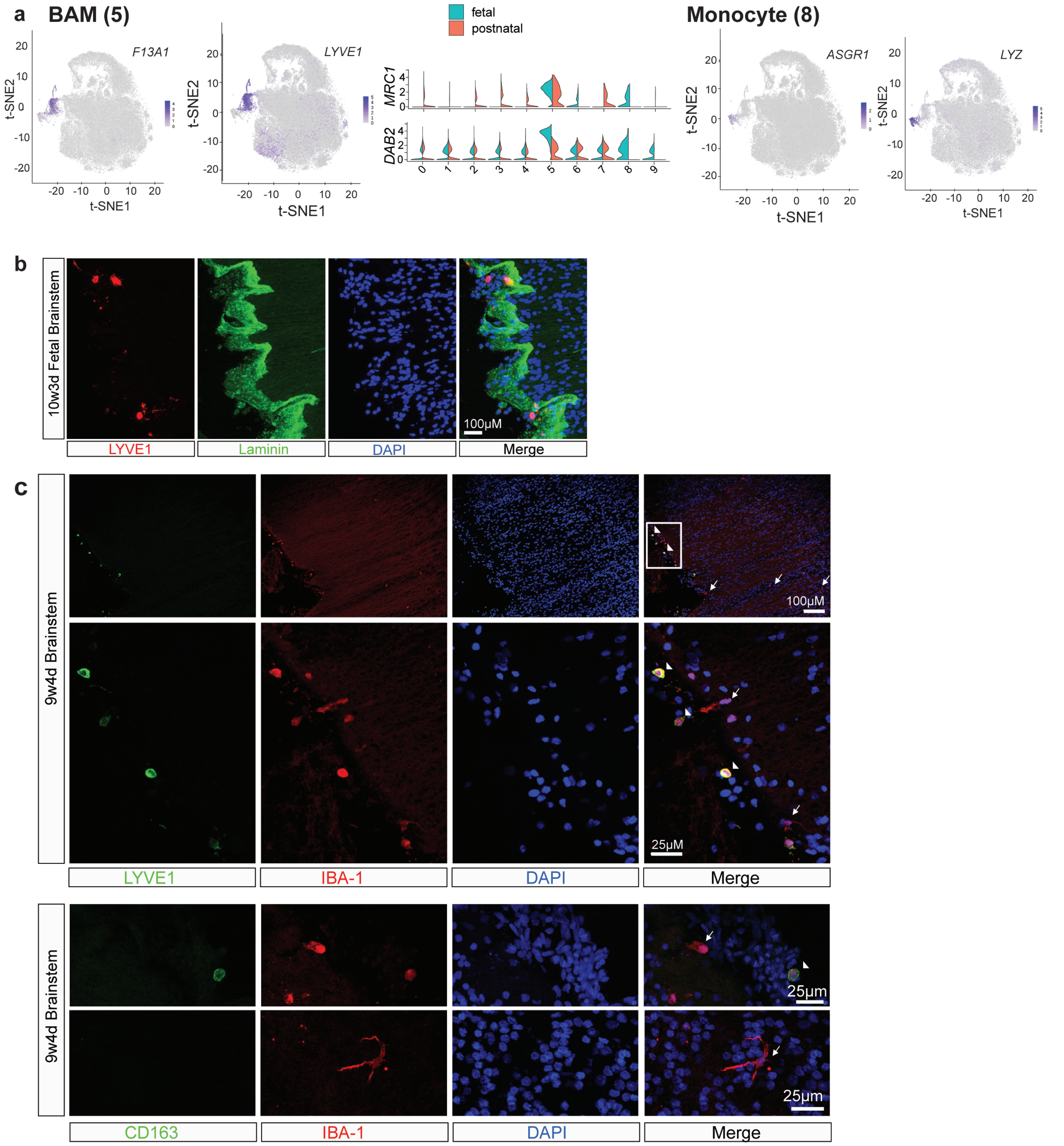

#### Extended Data Figure 8

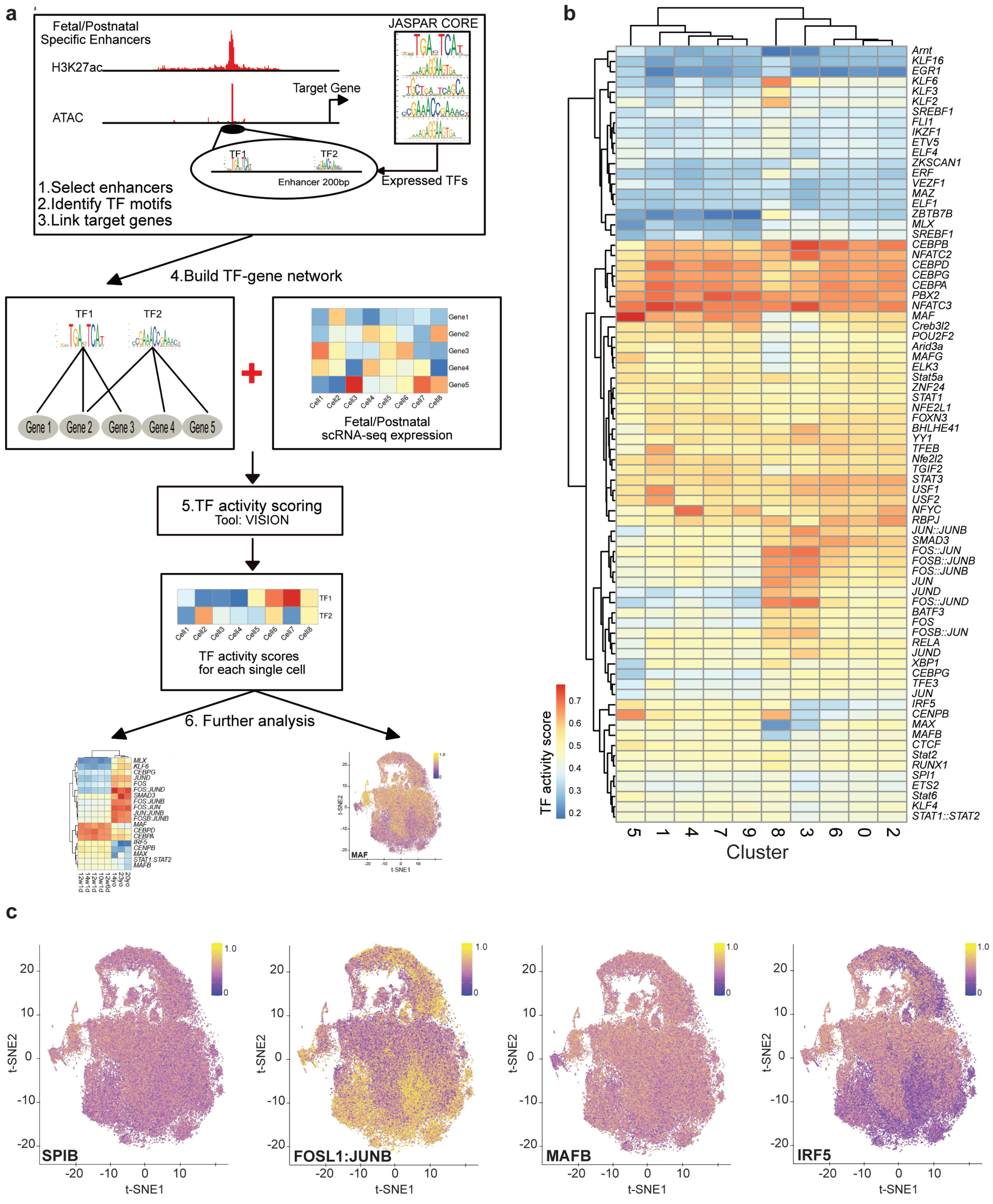

#### Extended Data Figure 9

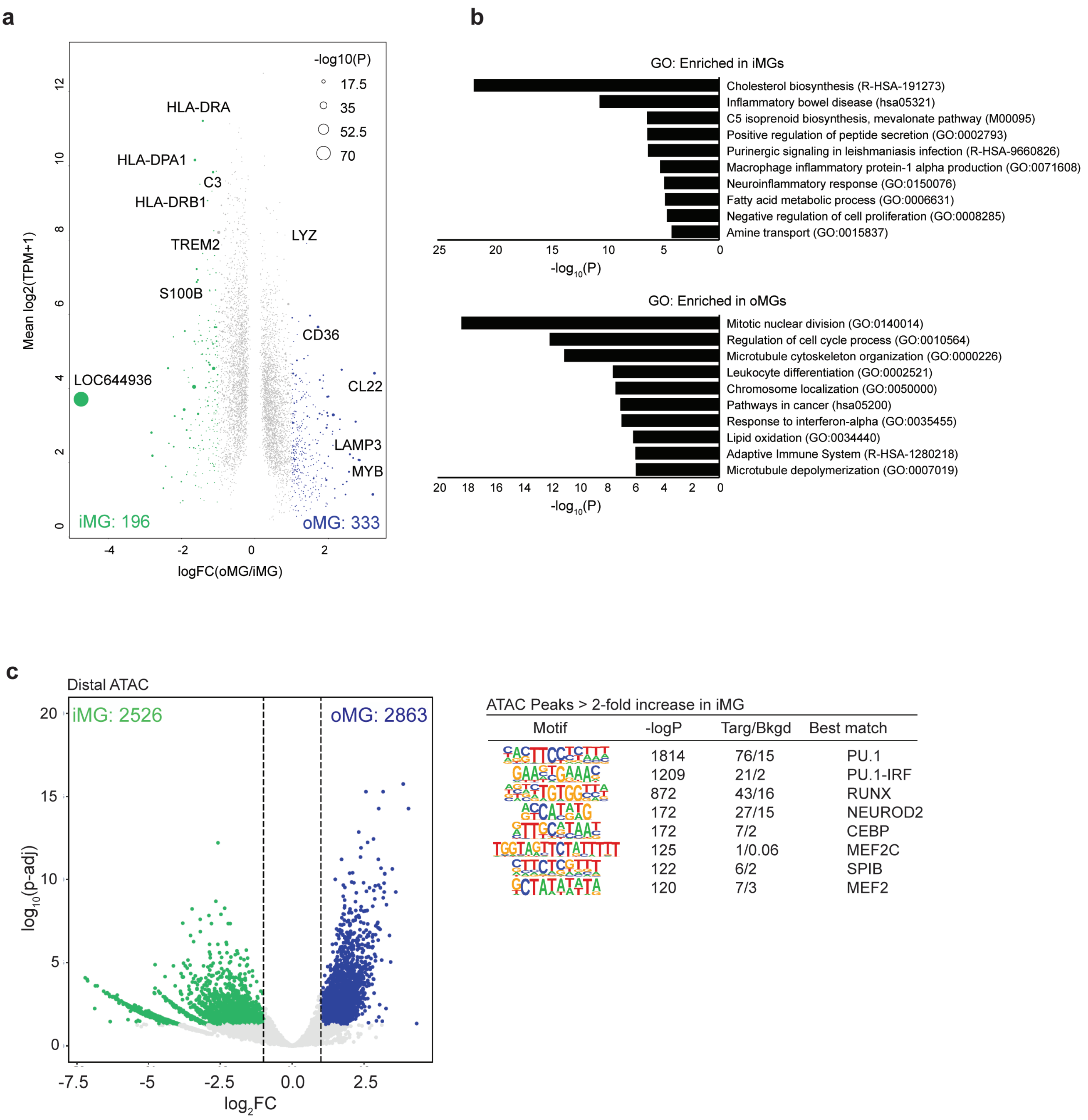

### Extended Data Figure 10

**a**

#### Lysosomal Storage Diseases

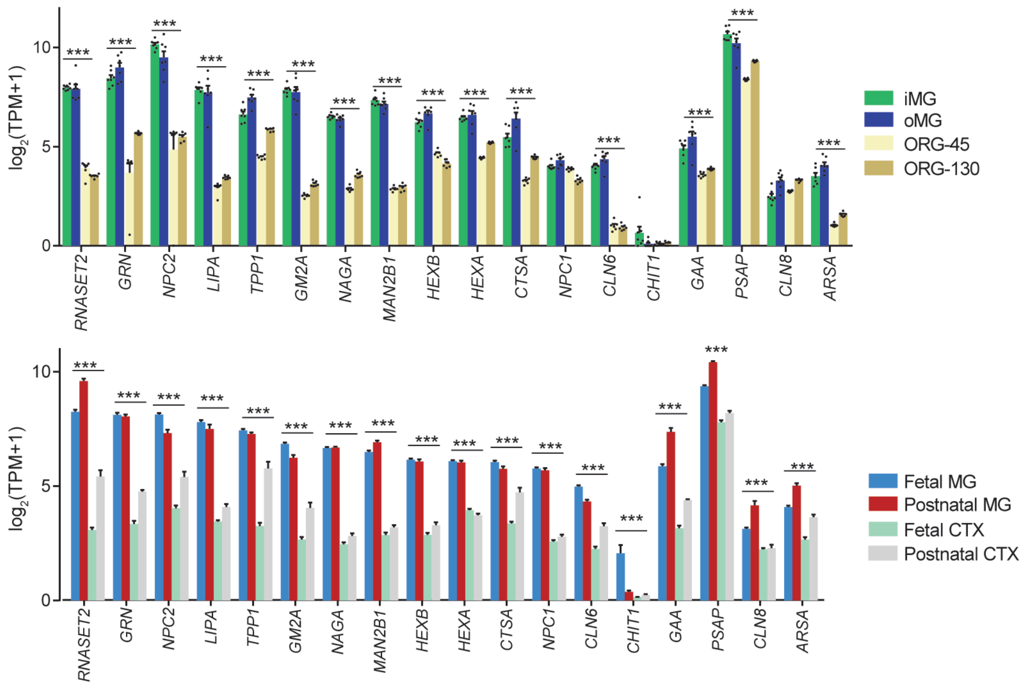


**b**

#### NPC phagocytosis

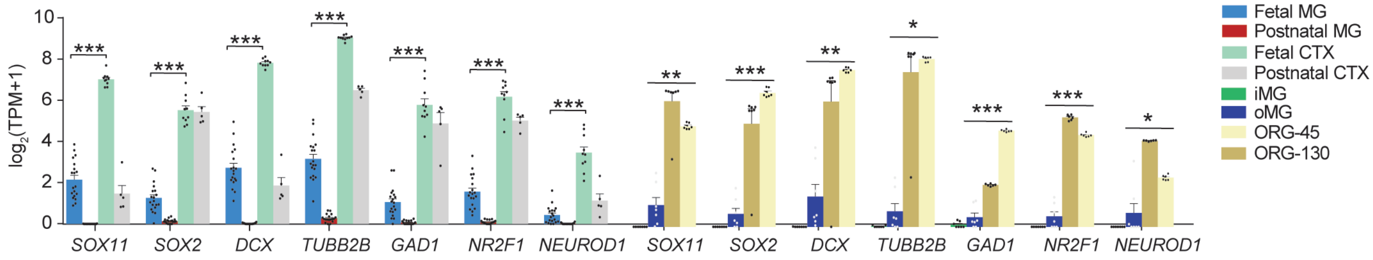


**c**

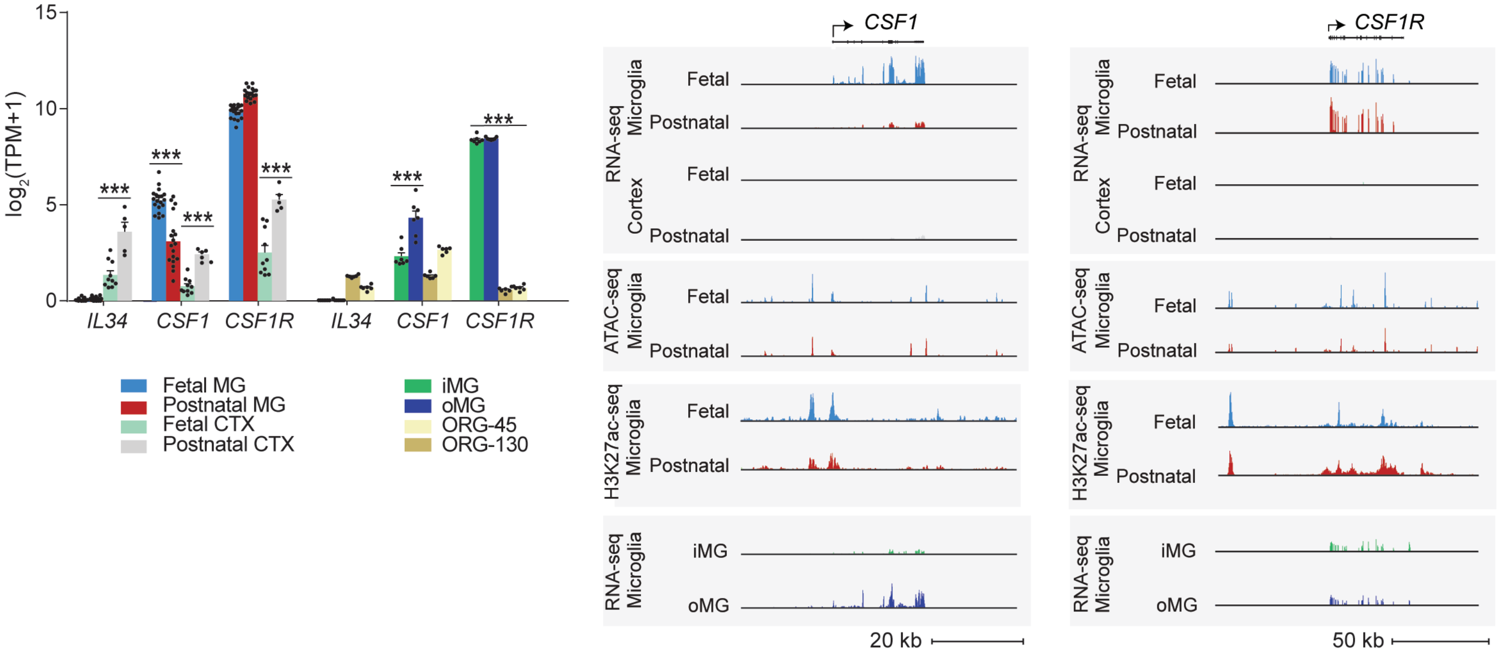
